# Homologues of key circadian clock genes present in *Verticillium dahliae* do not direct circadian programs of development or mRNA abundance

**DOI:** 10.1101/2019.12.20.883116

**Authors:** Emma Cascant-Lopez, Susan K. Crosthwaite, Louise J. Johnson, Richard J. Harrison

## Abstract

Many organisms harbour circadian clocks that promote their adaptation to the rhythmic environment. While a broad knowledge of the molecular mechanism of circadian clocks has been gained through the fungal model *Neurospora crassa*, little is known about circadian clocks in other fungi. *N. crassa* belongs to the same class as many important plant pathogens including the vascular wilt fungus *Verticillium dahliae.* We identified homologues of *N. crassa* clock proteins in *V. dahliae,* which showed high conservation in key protein domains. However, no evidence for an endogenous, free-running and entrainable rhythm was observed in the daily formation of conidia and microsclerotia. In *N. crassa* the *frequency* (*frq*) gene encodes a central clock protein expressed rhythmically and in response to light. In contrast, expression of *Vdfrq* is not light-regulated. Temporal gene expression profiling over 48 hours in constant darkness and temperature revealed no circadian expression of key clock genes. Furthermore, RNA-seq over a 24 h time-course revealed no robust oscillations of RNA in constant darkness. Comparison of gene expression between wild-type *V. dahliae* and a *ΔVdfrq* mutant showed that genes involved in metabolism, transport and redox processes are mis-regulated in the absence of *Vdfrq*. In addition, *VdΔfrq* mutants display growth defects and reduced pathogenicity in a strain dependent manner. Our data indicate that if a circadian clock exists in Verticillium, it is based on alternative mechanisms such as post-transcriptional interactions of VdFRQ and the WC proteins or the components of a FRQ-less oscillator. Alternatively, it could be that whilst the original functions of the clock proteins have been maintained, in this species the interactions that generate rhythmicity have been lost or are only triggered when specific environmental conditions are met. The presence of conserved clock genes in genomes should not be taken as definitive evidence of circadian function.

**Author summary:** Circadian clocks are used by organisms to orchestrate the activity of cellular processes such that they occur at an optimal time of day. Research carried out in the filamentous fungus *Neurospora crassa* has revealed a huge amount of information about the components its circadian clock, its interactions with the environment and how it drives cellular biochemistry and physiology. Although homologues of the Neurospora clock genes are present in a number of fungi, functional clocks have been demonstrated in a just a handful. Importantly, a link between the circadian clock of the plant pathogen *Botrytis cinerea* and virulence has recently been reported. We report that another significant plant pathogen, *Verticillium dahliae,* contains well-conserved homologues of all key clock genes. We find that diurnal development of conidia and microsclerotia is not influenced by a circadian clock. Furthermore, in a constant environment we find no evidence of rhythmic transcript accumulation. However, deletion of the central clock component results in altered growth and reduced virulence. This led us to question the role of clock genes in Verticillium. We are forced to consider that in this species the interactions that generate rhythmicity have been lost, are generated purely via post-transcriptional modification of clock proteins, are only triggered when specific environmental conditions are met or never evolved.

## Introduction

Circadian clocks are endogenous timekeepers that enable organisms to anticipate cyclic changes in the environment and thus confer adaptive advantage [1–2]. The defining characteristics of circadian clocks are: rhythmicity that persists in constant conditions (absent cyclic conditions) with a period of approximately 24 hours; the ability to entrain to external signals such as light and temperature; and temperature and nutritional compensation of period [3]. Such oscillations have been widely observed in most branches of life, and are particularly well characterized in *Neurospora crassa*.

In *N. crassa*, the circadian clock is based on a transcription-translation negative feedback loop [4], initiated by the photoreceptor and transcription factor White Collar-1 (WC-1) [5]. In the dark, WC-1 and White Collar-2 (WC-2) dimerize through their PAS domains, forming the White-Collar Complex (WCC) [6]. The WCC binds to the clock-box motif in the promoter of the *frequency* (*frq*) gene, activating its transcription [7]. When FRQ is synthesized, it associates with FRQ-interacting RNA helicase (FRH) forming the FCC complex. FCC inhibits the activity of the WCC, and consequently transcription of *frq* is reduced. FRQ is progressively phosphorylated by the kinase CK1, and degraded by the FWD-1 protein [8]. This leads to a reduced FCC-mediated inhibition of the WCC, which results in initiation of a new cycle [1]. In addition to the FRQ/WC-based oscillator (FWO) described above, *N. crassa* contains other FRQ-independent oscillators known as FLOs [9–11]. For example, the activity of nitrate reductase oscillates in the absence of FRQ [12], and a cryptochrome-dependent oscillator (CDO) [13] has also been described.

Both light and temperature can entrain the clock. Light rapidly induces transcription of *frq*, which leads to clock resetting and entrainment to the external stimulus [14]. The WCC complex activates *frq* transcription by binding to light responsive elements (LREs) present in the *frq* promoter [7]. Two LREs have been identified in the *frq* promoter; the proximal LRE is necessary for light induction of *frq,* whereas the distal LRE (known as the Clock-Box) is required for both light and circadian clock regulation [15–16]. Additionally, the WCC is essential for the transcriptional activation of numerous light-regulated genes, including other transcription factors, leading to a genetic regulatory cascade [17]. A subset of the downstream genes are known to be regulated by both the circadian clock and light. Included in this subset is the photoreceptor VVD, which is involved in photo-adaptation [18]. Despite not being essential for circadian function, VVD enhances the robustness of the clock by preventing clock resetting at dawn and promoting clock resetting at dusk [11,17–20]. Other targets include genes that function in carotenoid synthesis and spore development [15]. Importantly, rhythmic mRNA levels in *N. crassa* are not only controlled by light and the circadian clock at the promoter level, but through post-transcriptional regulation [21]. In *N. crassa* the circadian clock regulates approximately 40 % of the genome and a quarter of expressed proteins [21–22].

Temperature compensation and temperature resetting of the clock is mediated post-transcriptionally [23]. *frq* transcript levels remain the same at different temperatures but FRQ protein cycles remain at a higher mean level at high temperatures [24]. VVD is also temperature regulated, and plays a role in temperature compensation of the clock [25]. In addition, the existence of a temperature-compensated FLO that requires the components of the WCC has been described [10, 25]. An example of a temperature-compensated FLO-output is the clock-controlled gene *ccg*-*16*, which has been demonstrated to oscillate rhythmically in the absence of FRQ [10].

Despite detailed knowledge of the *N. crassa* circadian clock, little is known about the existence of functional circadian clocks and their molecular basis in other fungi, although homologues of the *N. crassa* core circadian clock proteins, especially WC-1 and WC-2, have been found in other fungal species [26–27]. It has been hypothesized that FRH and FWD-1 were present in the common ancestor of all fungi [28]. WC-1 and WC-2 were probably gained in the common ancestor of Zygomycetes, Basidiomycetes and Ascomycetes and subsequently lost in the Saccharomycetes. FRQ was likely gained in the ancestor of Ascomycetes and lost in Eurotiomycetes, remaining in Sordariomycetes, Leotiomycetes and Dothideomycetes [28–29]. Interestingly, FRQ is the least conserved of the clock proteins. While examples of functional *frq*-dependent circadian oscillators are present in the Leotiomycete *Botrytis cinerea* [30] and in the Pezizomycete *Pyronema confluens* [29], other fungi, such as the Dothideomycete *Aureobasidium pullulans,* show no rhythmic *frq* expression, although they do display a circadian developmental rhythm [31]. Additional circadian rhythms have been demonstrated in species lacking a *frq* homologue, such as in *Aspergillus flavus* and *Aspergillus nidulans* [32]. Furthermore, the yeast *Saccharomyces cerevisiae* was shown to exhibit circadian entrainment of metabolism despite the absence of a *frq* homologue [33], indicating a widespread presence of circadian oscillatory molecular processes across fungal species.

*N. crassa* is phylogenetically related to many important plant pathogens including *Verticillium dahliae*. *V. dahliae* is an asexual soil-borne Sordariomycete that causes wilt disease on more than 200 plant species worldwide, including high-value agricultural crops [34]. Verticillium sp. can persist in soil for long periods in the form of melanized resting bodies. In *V. dahliae*, these are clusters of thick-walled, septate and dark pigmented hyphal cells called microsclerotia [35]. Microsclerotia germinate upon the detection of plant root exudates, and colonize the vascular system. Once inside the vessels, conidia are produced and transported upwards reaching extensive portions of the plant. Blockage of the transport system results in wilting of the plant and the subsequent formation of microsclerotia in the dead tissue, concluding the disease cycle.

The aim of this work was to determine whether a circadian clock is present in *V. dahliae*. As previously described [36], under 24 hour light/dark cycles *V. dahliae* displays concentric rings of conidia and microsclerotia. We report that this phenotype is directly driven by external cues, rather than entrained. Comparative genetic studies between *N. crassa* and *V. dahliae* reveal that key clock components are conserved not only at the domain level, but down to individual phosphorylation sites. Similar sequences to the proximal and distal LRE motifs are present upstream of the VdFRQ ORF, although they are very widely spaced. These binding motifs may be important for a WC-mediated light-response. However, qRT-PCR studies over a 48-hour time-course revealed no rhythmic *Vdfrq* expression under constant conditions or in cycles of light/dark and high/low temperature. RNA-sequencing gene expression studies in the WT revealed large-scale changes in gene expression driven by changes from light to dark, but time-course RNA-sequencing revealed no strong signature of gene expression indicative of circadian rhythmicity. Our results show that while there is high conservation of clock components between *V. dahliae* and *N. crassa,* there is no strong evidence of a functional circadian clock in *V. dahliae,* at either the physiological or the molecular level.

## Methods

### Identification of *V. dahliae* putative clock-genes

*N. crassa* OR74A and *V. dahliae* JR2 genomes were downloaded from Ensembl Fungi [37]. Homologues were identified using BLASTp against the *V. dahliae* JR2 genome. Whole gene alignments between the query *N. crassa* gene and the *V. dahliae* JR2 hit were carried out in Geneious R10 using the MUSCLE algorithm. Next, *V. dahliae* JR2 hits were aligned to the genomes of five *V. dahliae* strains isolated and sequenced at NIAB EMR (12253, 12158, 12251, 12161 and 12008), available at DDBJ/EMBL/GenBank database using the following numbers: PRJNA344737 and PRJNA352681. Domains were identified within predicted proteins using InterProScan. Nuclear localization signals (NLS) were identified using cNLS Mapper. Identification of clock gene homologues in *V. albo-atrum* (PD747)*, V. alfalfae* (PD683)*, V. nonalfalfae* (TAB2)*, V. longisporum* subgenome A (VLB2)*, V. longisporum* subgenome D*, V. nubilum* (PD621)*, V. tricorpus* (PD593)*, V. isaacii* (PD660)*, V. klebahnii* (PD401) and *V. zaregamsianum* (PD739) was performed using BLASTp in collaboration with the Thomma Lab (Wageningen University).

### Orthology analysis

The study involved 25 species, mostly plant pathogens, sampled from across the phylum Fungi. The respective genome (repeat-masked), CDS, and protein sequences were downloaded from Ensembl Fungi [37]. The orthology relationships between the genes in each genome were first established using the OrthoFinder ver. 1.0.7 [38] and OrthoMCL ver. 2.0.9 [39] pipelines. The genomes of 25 fungal species were searched for homologues of *frq* (NCBI ID: 3876095)*, wc-1* (NCBI ID: 3875924)*, wc-2* (NCBI ID: 3879968), *frh* (NCBI ID: 3872445)*, fwd-1* (NCBI ID: 3872130) and *vvd* (NCBI ID: 3873728). The phylogenetic tree was built using a concatenated alignment of DI/D2 regions of large subunit (LSU) rDNA and ITS regions that were identified through BLAST and extracted from the genomes. The Tamura-Nei method [40] was used to build trees, as implemented within Geneious R10.

### Promoter Motif Identification

2000 bp upstream and downstream of each gene ORF were extracted. MEME suite ver. 4.11 was subsequently used to analyse the motif content of promoter regions [41]. For motif scanning purposes, the program FIMO [42] was used to deal with ungapped motifs (ACE motif in *ccg2* promoter), while gapped motif search for sequences showing similarity to the Clock-box element in *frequency* promoters was carried out with GLAM2SCAN [43]. For *de novo* motif discovery, DREME [44] and GLAM2 were used for ungapped and gapped motifs, respectively. Motif enrichment analysis was carried out with the AME program. A manual screen of consensus LRE motifs described in [15, 45] on the 2000 bp upstream and 2000 bp downstream regions of the *frq* orthologues across 13-species was performed using Geneious R10.

### *V. dahliae* isolates and growth conditions

Strains of *V. dahliae* (**S1 Table**) were stored at −80 °C as conidial suspensions preserved in 50 % glycerol. Isolates were cultivated on petri dishes containing Prune Lactose Yeast Agar (PLYA) [46] at 24 °C in constant darkness. After a week, plates were flooded with 2 ml sterile water and gently rubbed using a plastic spreader to create a spore suspension.

### Plate assays

Conidial suspensions were filtered using filter paper with 3 to 5μm pore size and 1 µL of a 3 x 10^5^ spores/mL suspension was point-inoculated in the centre of a PLYA plate and incubated for 14 days under the appropriate light and/or temperature conditions. For light entrainment experiments, after 14 days at 24 °C in 12:12 LD, plates were marked at the end of the dark cycle before transfer to constant darkness (DD) for 7 days. For temperature entrainment assays, the cultures were incubated under 12 h at 20 °C followed by 12 h at 28 °C in the dark during 14 days prior to transfer to constant temperature. To assess the effect of entrainment by both light and temperature, plates were incubated in 12 h light at 28 °C/12 h dark at 20 °C, before transfer to constant darkness and temperature conditions. To examine nutritional compensation cultures were inoculated onto a high-nitrogen medium (PDA), low-nitrogen medium (Czapek DOX Agar), minimal medium (MM) [47] and basal minimal medium (BMM) [48] and incubated in 12:12 LD cycles for 14 days. Strains were incubated in Panasonic MLR-352 or MIR-154 incubators equipped with broad spectrum fluorescent lamps FL40SSENW37. The light intensity was 75 µmol s^-1^ m^-2^ (approximately 5600 lux). Colony size was measured using an electronic calliper. Each experiment was repeated at least three times and contained three replicas of each strain and treatment.

### Time-course and light pulse experiments

Rhythmic expression analysis by qRT-PCR in free-running conditions (constant darkness and constant temperature) was similar to the protocol described in [49]. A mycelial mat was formed by inoculating 1 x 10^8^ spores of *V. dahliae* into Petri dishes containing 20 ml of half strength PDB medium and incubated at 25 °C under constant light for 96 hours. Individual mycelial disks (1 cm diameter) of mycelium were cut and inoculated in 100 ml Erlenmeyer flasks containing 25 ml of half strength PDB medium. A total of 36 flasks were used to cover 12 sequential time-points throughout two circadian cycles (4 h, 8 h, 12 h, 16 h, 20 h, 24 h, 28 h, 32 h, 36 h, 40 h, 44 h, 48 h), each replicated three times. The flasks were incubated at 25 °C in constant light for at least 24 hours under agitation at 120 rpm. Cultures were staggered into constant darkness at different intervals, so that when harvested cultures were of similar ages. Cultures were harvested under red light. Disks were dried with filter paper, placed into 2 ml Eppendorf tubes, frozen in liquid nitrogen and stored at −80 °C.

A similar methodology to that described in [50] was performed for light and temperature entrainment analysis. A total of 36 flasks were used to cover 12 sequential time-points throughout one LD cycle (L2, L4, L6, L8, L10, L11.5, D2, D4, D6, D8, D10, D11.5). Similarly, a total of 36 flasks were used to cover 12 sequential time-points throughout one cycle of 12 h at 20 °C (low temperature, LT) /12 h at 28 °C (high temperature, HT) (L2, L4, L6, L8, L10, L11.5, H2, H4, H6, H8, H10, H11.5).

For light pulse experiments, flasks containing half strength PDB medium were inoculated with *V. dahliae* or *N. crassa* mycelial discs and subsequently grown in constant light for 24 hours at 25 °C (120 rpm), after which they were transferred to constant darkness for 30 hours prior to a 15 minutes exposure to white light (75 µmol s^-1^ m^-2^). After the light pulse, cultures were harvested under red light. Control flasks were not exposed to light pulses and were harvested under red safe-light conditions.

### RNA extraction

Frozen mycelium of *N. crassa* and *V. dahliae* were ground with a mortar and pestle and total RNA extracted using the RNeasy Plant Mini Kit (Qiagen GmbH, Germany) according to the manufacturer’s instructions. The integrity of RNA samples was assessed using the Agilent 4200 TapeStation system (Agilent, Santa Clara, USA).

### Gene expression analysis by Quantitative Real Time PCR (qRT-PCR)

cDNA from 1 µg of RNA was synthesized using the QuantiTect Reverse Transcription Kit (Qiagen, Germany) following the manufacturer’s instructions. qRT-PCR was performed with SYBR green (qPCRBIO SyGreen Mix Lo-Rox, PCR Biosystems, UK) and amplification followed using the CFX96™ Real-Time PCR detection system (Bio-rad). Normalisation was carried out against *V. dahliae* β-tubulin and Elongation factor 1-⍺ transcripts. For *N. crassa*, *β-tubulin* and TATA binding box-encoding gene served as housekeeping genes. Primers are listed in **S2 Table**. PCR reactions were carried out in 10 µl containing 400 nM of each primer, 5 µl of 2x qPCRBIO SyGreen Mix and 2 µL of 1:4 diluted cDNA sample. The PCR conditions were 95 °C 2 min; 39 cycles of 95 °C 10 sec, 60 °C 10 sec, 72 °C 30 sec and 1 cycle of 95 °C 10 sec, 65 °C 5 sec 95 °C 5 sec. Relative gene expression was calculated using the comparative cycle threshold (C_T_) method (2^-(ΔΔCt)^) method [51]. Expression values for free running time-course experiments were normalized to the cultures grown 4 h in the dark (D4). Expression values for time-course experiments under LD cycles were normalized to the cultures grown 2 h in the dark (D2), and cultures grown 2 h at 20 °C (2L) were the reference samples for the temperature cycle/entrainment time-courses. The expression values for light pulsed cultures were normalized to cultures maintained in control conditions (DD).

Expression values derived from three biological replicates (unless otherwise stated) each containing three technical repeats, were analysed using one-way Analysis of Variance (ANOVA). The residuals were tested for normality and if required, data were log-transformed. Statistical analyses were carried out using R Studio software. Statistical analysis of rhythmicity of time-course experiments was achieved with JTK-CYCLE [52] in the R software (version 3.3.0) using the 2^-(ΔΔCt)^ normalized data of all the replicates. The analysis was performed as described in the JTK-CYCLE manual, looking for period lengths of 24 h.

### RNA-Seq

For the time-course RNA-seq experiment in free-running conditions, samples of *V. dahliae* 12253, 12008 and *ΔVdfrq_12253* were harvested under red light after 6 h, 12 h, 18 h and 24 h in the dark. Concurrently, for light versus dark analysis, cultures were incubated in constant darkness and then were transferred to white-light and harvested after 6 hours. The experiment contained three biological replicates of each strain and condition.

Total RNA was extracted as described above. For transcriptome sequencing, samples with a minimum of 1 µg of RNA (100 ng/µl), ≥ 6.8 RIN and 260/280 nm values > 1.8 were sent to Novogene Technology Co. Ltd. (Wan Chai, Hong Kong). Sequencing was performed on Illumina HiSeq P150.

Quality control of the RNA-seq reads was carried out by fastQC and adapters were trimmed using fastq-MCF (available from https://expressionanalysis.github.io/ea-utils/). STAR software [53] was used to align RNA-seq reads to the reference *Verticillium dahliae* JR2 genome (Ensembl Fungi). Mapped read counts were calculated using the program featureCounts [54]. The analysis of expression of all predicted genes was performed with the DESeq2 package in R. A grouping command was used to assess differentially expressed genes (DEG) between conditions (L or D) and strains (Vd12253, Vd12008 and *ΔVdfrq_12253)*. The false discovery rate (FDR) cut-off was set to 0.05. Genes were considered to be significantly differentially expressed when *p*-value < 0.05 and presented more than 1-log2 fold change (LFC) in transcript level.

For rhythmic expression analysis, the raw counts of the time-course samples were normalized to fragments per kilobase of transcript per million mapped fragments (FPKM). The time-course of each individual strain was analyzed using JTK-CYCLE, looking for rhythms of 24 h [52]. Genes with a *p*-value < 0.05 and a false discovery rate of q < 0.1 were considered rhythmic.

Principal component analysis (PCA) plot and samples distances plot used rlog-transformed read counts, and were carried out using R. Gene ontology (GO) terms were retrieved from GO.db in R. Gene enrichment analysis for GO terms was performed using topGO in R with Fisher’s exact test to retrieve significantly enriched processes of the DEG. The analysis of secondary metabolite biosynthetic gene clusters in *V. dahliae* was performed using antiSMASH [55].

### Construction of *V. dahliae frq* replacement cassette

The 5’ and 3’ flanking regions of the *Vdfrq l*ocus were amplified from genomic DNA using primer pairs HRFrq1-F/R and HRFrq2-F/R respectively (**S2 Table**). Core USER Bricks and the *hygromycin* resistance cassette were amplified from pRF-HU2 plasmid and the vector bricks were assembled through USER reaction and transformed into *E. coli* DH5α competent cells [56]. The resulting vector junctions were amplified using B1.B2-F/R, B1.B2.F1.H-F/R and H.F2.B1-R to confirm the correct assembly.

### Agrobacterium tumefaciens transformation

*Agrobacterium tumefaciens* EHA105 was transformed with the *Vdfrq* replacement vector as described in [57]. Putative transformants were checked by PCR using primers B1.B2-F/R, B1.B2.F1.H-F/R and H.F2.B1-R (**S2 Table**).

### Agrobacterium-mediated transformation of *V. dahliae*

*A. tumefaciens* EHA105 containing the pEcFrq-D1 binary vector was grown at 28 °C for 48 h in MM containing 50 μg ml^-1^ of rifampicin and 50 μg ml^-1^ of kanamycin on a rotary shaker (200 rpm). At an optical density of OD_600_ = 0.5, bacterial cells were diluted to OD_600_ = 0.15 with IM [58] containing 200 μM acetosyringone (AS). Cells were grown for an additional period of 12-15 h before being mixed with an equal volume of a spore suspension (1×10^6^ spores ml^-1^) of the required strain of *V. dahliae*, previously grown for three days in flasks containing PDB liquid media in a rotary shaker 180 rpm in the dark. From this mixture aliquots of 200 μl were plated on a cellophane membrane placed on an IM agar plate. After incubation at 25 °C for 48 h the membrane was transferred onto a PDA plate containing 200 μg/ml tricarcillin and 50 μg/ml hygromicin B and incubated at 24 °C for a further 5 to 7 days. Discrete colonies were selected and grown for 24 h in PDB liquid cultures containing 50 μg/ml hygromycin B. Single-spore colonies were obtained by plating 100 μL of the culture on PDA with 50 μg/ml hygromycin B. PCR was carried out on selected single-spore mutants to confirm insertion of hygromicin using the primers TestHygr-F/R and FrqUS_Hygr-F/R, and lack of product from deleted gene using the primers TestFrq-F/R **(S1 Fig.)**. The PCR conditions using Taq 5x Master Mix (New England Biolab) were 95 °C 30 sec; 35 cycles of 95 °C 20 sec, 60 °C 20 sec, 68 °C 2 min and 68 °C for 5 min.

### *Arabidopsis thaliana* and strawberry *in vitro* propagation

For pathogenicity tests, seeds of *Arabidopsis thaliana* ecotype Columbia (Col), were surface-sterilized by sequential immersion in 70% ethanol for 1 min and 10% (v/v) bleach containing 0.1% (v/v) Tween-20 for 5 minutes with gentle shaking. Seeds were then washed with autoclaved distilled water for 5 minutes five times, re-suspended in sterile 0.1% agarose, and subsequently stratified at 4 °C over a period of 2-3 days in darkness [59]. 5 to 6 seeds were pipetted into 120 x 120 mm square petri dishes (Thermo Fisher Scientific) containing half-strength Murashige and Skoog salts (MS), pH 5.7; 0.8% (wt/vol) Phytagel (Sigma-Aldrich); 1% sucrose; and grown at 22°C under 16:8 LD cycles in a Panasonic MLR-352 incubator for 3-4 weeks.

Strawberry (*Fragaria x ananassa*) cultivar Hapil plantlets were preserved in sterile jars containing with SMT proliferation medium (MS salts 4.41 g, 177.5 μM Benzylamino purine (BAP), 254 μM Gibberellic acid (GA), 255 μM Indole butyric acid (IBA), sucrose 30 g, agar (Oxoid 3) 7.5 g, adjust pH = 5.6). The optimal growth conditions were 24 °C in the light and 19 °C in the dark, in a photoperiod of 16 h light and 8 h dark. For long-term in vitro material, plantlets were transferred to fresh SMT medium in sterile conditions every 6 weeks. For root induction, SMR rooting medium (MS salts 4.41 g, 254 μM GA, 255 μM IBA, sucrose 30 g, agar (Oxoid 3) 7.5 g, adjust pH = 5.6) was used. Four weeks prior to a pathogenicity test, roots were carefully removed from the media just below the crown and the tips were moved to square petri dishes containing ATS medium as specified in [60], where subsequent roots were produced.

### *Arabidopsis thaliana* and strawberry root inoculation assay

*V. dahliae* conidial suspensions were harvested from 7 day old liquid cultures and diluted to 1 x 10^6^ spores/ml with PDB. Three to four-week-old *Arabidopsis* seedlings were moved under aseptic conditions to new plates containing half-strength MS salts pH 5.7; 0.8% (wt/vol) Phytagel (Sigma-Aldrich) without sucrose, and root tips were trimmed using sterilized scissors. Each plate contained five *Arabidopsis* seedlings that were inoculated with 200 μL of the spore suspension spread onto the roots. In mock plates seedlings were inoculated with 200 μL of autoclaved distilled water. Plates were subjected to a light-dark cycle of 12:12 h and constant temperature of 24 °C for the duration of the experiment.

For the strawberry inoculation, plants were removed from the plates and the roots were carefully cleaned free of agar with a sterile blade. Before the inoculation, 1 cm of roots was trimmed from the bottom with sterile scissors. Plants were inoculated by dipping the root systems into a flask containing 1 x 10^6^ spores/ml suspension (6 plants/100 ml inoculum) for 5 minutes. A fresh spore suspension was used for each batch of 6 plants. Plants were then placed to new ATS medium plates sealed with tape. Uninfected plants were inoculated with sterile distilled water following the same procedure as stated before. Plates were subjected to a light-dark cycle of 12:12 h and temperature of 24 oC in a Panasonic MLR-352 incubator.

### Disease assessment

*Arabidopsis* and strawberry cv. Hapil seedlings were photographed and scored for disease symptoms using a scale with nine categories modified from [61]; Score 1: Healthy plant, 2: Slight symptoms on older leaves (yellowing), 3: 1-2 outer leaves affected, 4: >2 leaves affected, 5: 50% leaves affected, 6:> 50% leaves affected, 7: 75% leaves affected, 8: >75% leaves affected, 9: Plant dead.

## Results

### Distribution of core clock gene homologues in Verticillium species and other Sordariomycete, Dothideomycetes, and Leotiomycetes of economic importance

The genomes of 25 Sordariomycete, Dothideomycete and Leotiomycete species were searched for homologues of *frq, wc-1, wc-2, frh, fwd-1* and *vvd*. The predicted proteins from these genomes were clustered and proteins in the same orthogroup cluster as the *N. crassa* genes were considered orthologous genes. Orthologues of all six clock oscillator genes are present in most species of Sordariomycetes tested in this study, including important plant-pathogenic fungi such as *Glomerella graminicola and Neonectria ditissima* (**Fig 1A**). Homologues of clock genes are present in all *Verticillium* species tested: *V. albo-atrum, V. alfalfae, V. nonalfalfae, V. dahliae, V. nubilum, V, tricorpus, V, isaacii, V. klebahnii* and *V. zaregamsianum* (pers. comm. B. Thomma). *Fusarium oxysporum* has three copies of *frq* and *vvd*. Conversely, members of the chaetomium family appear to have lost *frq, wc-1,* and *vvd* homologues. In contrast to *Sclerotinia sclerotiorum* and *Botrytis cinerea*, the Leotiomycete *Blumeria graminis* lacks a *vvd* homologue, but contains copies of the other clock components. *vvd* is absent in the Dothideomycetes *Cercospora zeae-maydis, Alternaria alternata* and *Venturia inaequalis,* but present in the wheat pathogen *Zymoseptoria tritici* that contains homologues of all the clock genes (**Fig 1A**). The genome of the Ustilaginomycetes *Ustilago maydis*, on the other hand, does not harbour an orthologue of *frq,* in agreement with the findings in [28].

**Figure 1.**
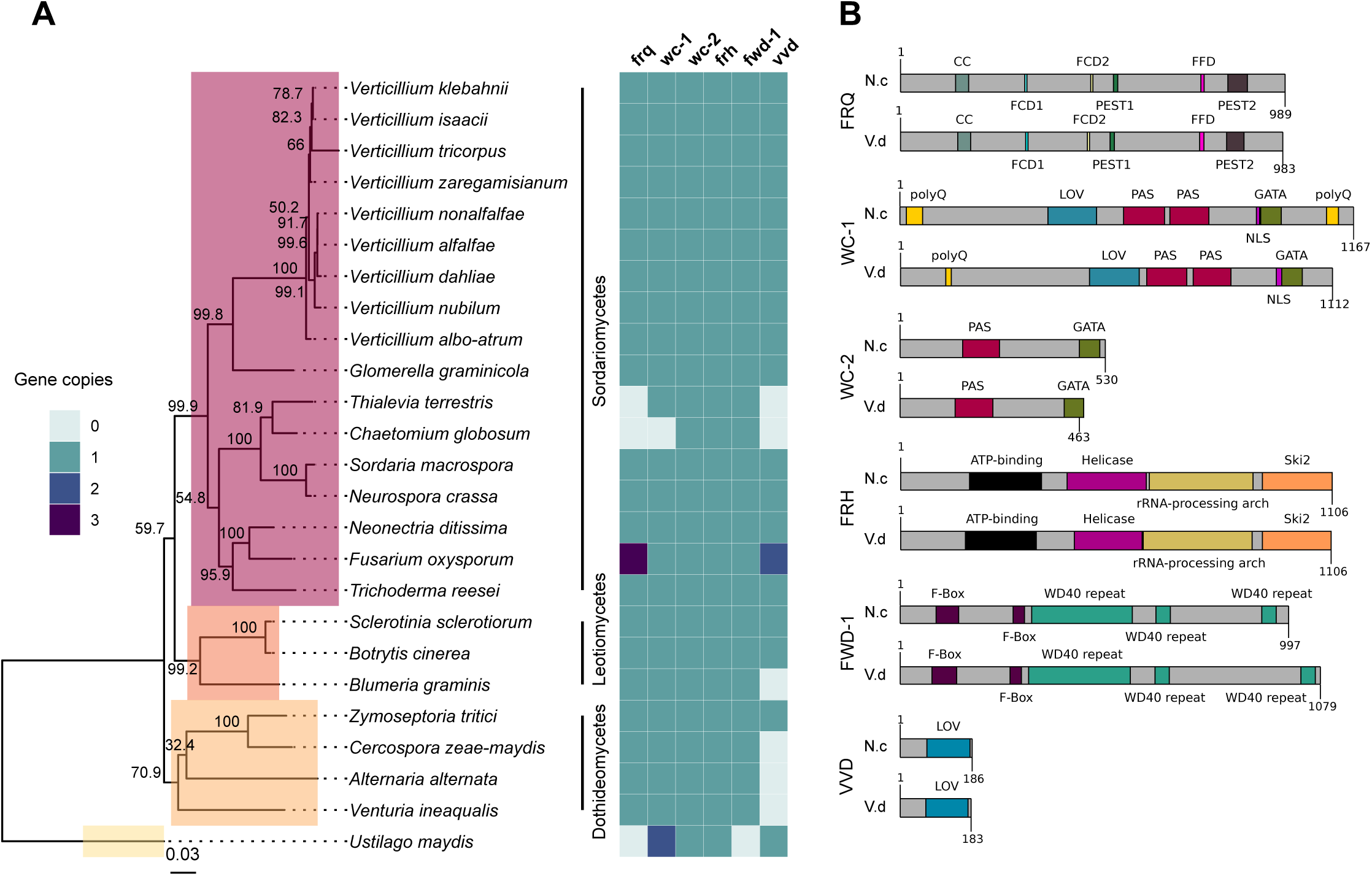
The circadian clock components are conserved across fungal species. **A.** *N. crassa* clock gene homologues across 25 fungal genomes. The presence, absence and copy number is shown for core clock genes (*frq, wc-1, wc-2, frh, fwd-1, vvd)* on the heatmap on the right. The phylogenetic tree was constructed based on the concatenated D1/D2 and ITS sequence alignment using the Neighbour-Joining method based on the Tamura-Nei model with Geneious R10 software. Bootstrap support values from 1000 replicates are shown at the nodes. The fungal classes are shown in color boxes on the tree. The Basidiomycetes Ustilago maydis was used as outgroup species. **B**. Protein domain structure of core clock proteins in *N. crassa* and *V. dahliae* based on InterProScan database. FRQ, WC-1, WC-2, FRH, FWD-1 and VVD are highly conserved at the domain level. Protein length is displayed at the end of each protein sequence. CC (coiled-coil), FCD and FRQ-CK1a (interaction domains), PEST (protein association), polyQ (poly-glutamine stretch domain), LOV (Light-oxygen-voltage sensing), PAS (Per-Arnt-Sim, protein binding domain), GATA (Zinc finger, GATA-type, DNA binding domain), Helicase (ATPase activity), F-box (protein-protein interaction motif), WD40 repeat (interacting domain).

Alignments of core clock proteins between *N. crassa* and *V. dahliae* revealed sequence identities greater than 43% and query coverages greater than 39%. Consistent with previous reports [29], *V. dahliae* FRQ is the least conserved (43.82% identity), and FRH is the best conserved (68.69% identity) of the core clock proteins. Clock proteins are strongly conserved at the domain level (**Fig 1B**). Although the FRQ protein alignment created between *N. crassa* and *V. dahliae* sequences shows moderate conservation overall, a number of important regions are highly conserved (**S2 Fig).** In *N. crassa* PEST1 and PEST2 are involved in determination of period length and cytoplasmic accumulation of the WCC [62–63]. Whilst a high degree of variation within VPEST2 exists, VPEST1 is better conserved. The formation of FRQ homodimers, essential for clock function, is mediated through its coiled-coil domain [64]. Between *N. crassa* and *V. dahliae* this domain shares 69.69% identity. FRQ-CK1a interaction domains, FCD1 and FCD2 [65–66] share 87.5% identity between *N. crassa* and *V. dahliae.* In contrast, low sequence conservation of the FRQ-FRH interaction domain (FFD) [67] is observed between *N. crassa* and *V. dahliae,* with 4 of 10 sites conserved. *V. dahliae* FRQ protein contains two predicted nuclear localization signal (NLS) sequences, one almost identical to the *N. crassa* NLS (DLLKRDKLFEIKVHGLPKPKKRELE). The NLS present in *N. crassa* FRQ is required for its entrance into the nucleus and down-regulation of *frq* transcription [68]. Furthermore, of the 73 *in vivo* and *in vitro* identified phosphorylation sites in *N. crassa* FRQ [69], 40 sites are conserved in *V. dahliae* FRQ. Of note Ser 513, a determinant of period length [70] and FRQ degradation, is present in *Vd*FRQ.

Alignment of *frq* homologues from six *V. dahliae* strains (JR2, 12253, 12251, 12008, 12161 and 12158) revealed several single nucleotide polymorphisms (SNPs) between isolates that are associated with their vegetative compatibility (VC) background (data not shown). For instance, associated with VC subclade II-2 (12161 and 12158) or VC subclade II-1 (JR2, 12253, 12251 and 12008). Consequently, VdFRQ protein alignment indicates four amino acid substitutions. However, none of these substitutions are in the known functional domains.

In *N. crassa*, WC-1 and WC-2 interact through their PAS domains to form a heterodimeric complex (WCC complex). The WCC is essential for light-induced gene expression of *frq* and other light-regulated genes, and is required for the generation of circadian rhythms [64, 71]. *V. dahliae* WC-1 and WC-2 contain all the domains required for their interaction, light perception and transcription factor (TF) activity (**Fig 1B**). *Vd*WC-1 and *Vd*WC-2 share 45.22% and 51.12% sequence identity with the *N. crassa* homologues, respectively. Nevertheless, *Vd*WC-1 is highly conserved (79.0% to 94.34% identity) in the known protein domains. *Vd* WC-1 contains a conserved Light-oxygen-voltage (LOV) domain. This domain is related to the Period-ARNT-Single-minded (PAS) domain family, and is distinguished by binding a flavin cofactor and the presence of a GXNCRFLQ motif [72]. Furthermore, VdWC-1 contains two highly conserved PAS domains required for protein interaction, and a GATA-type zinc finger DNA binding domain commonly present in GATA-type transcription factors (**S3 Fig**). A region of basic amino acids (LLSNKKKRKRRKGVG) required for rhythmicity and circadian expression of the Neurospora circadian clock gene *frequency* is also present [73]. Despite the high level of conservation of most WC-1 domains between *N. crassa* and *V. dahliae*, the conservation is poor in the first 480 aa. In *N. crassa*, this region contains a poly-glutamine (Poly-Q) region expansion aa 16-61 that has previously been observed in transcription factors and is reported to play a role in transcription activation and to affect period length [74–75]. Other reports suggest that neither the N nor C-terminal polyQ stretches are required for activation of *frq* transcription, and it is an adjacent region spanning aa 100-200 in WC-1 that is necessary for clock control of *frq* expression in the dark but not for light responsiveness of *frg* [76]. In *V. dahliae*, however, the polyQ stretch is not continuous and conservation is minimal in the aa 100-200 region. Furthermore, *V. dahliae* WC-1 lacks the C-terminal (Poly-Q) region present at the aa 1091-1133 in *N. crassa*.

*Vd*WC-2 contains a single PAS domain, a GATA-type zinc finger transcription factor domain with 65.71 % and 79.24 % sites conserved. A putative NLS (RGRKRKRQW) sequence identified in *V. dahliae*, lacks a homologous sequence in *N. crassa* (**S4 Fig**).

### *V. dahliae* conidiation and microsclerotia production are light-regulated but not under control of a circadian clock

*V. dahliae* produces visible concentric rings of conidia and microsclerotia when cultured in cyclic environments, such as light/dark (LD) and high/low temperature cycles [36]. Under 24 hour LD cycles at constant temperature, hyphal growth occurs under both light and darkness, however conidiation from hyphae is induced after a period of light (**Fig 2A and B**). After a minimum of 48 hours of growth microsclerotia form but in LD cycles they are mainly produced in the dark [36]. Nevertheless, in either constant darkness (DD) or constant light (LL) conditions *V. dahliae* cultures contain conidia and microsclerotia but they are not produced in a ring fashion (**Fig 2B**). To test if development is controlled by a circadian clock we investigated whether the morphological rhythms found in this pathogen are endogenous and can be entrained. We found that *V. dahliae* cultures transferred to DD after an entrainment period of 12:12 LD for 14 days at 22 °C cease to produce rings of conidia and microsclerotia. In 12:12 temperature cycles of 20 °C and 28 °C in DD or LL the banding pattern is synchronized to the temperature oscillations, and ceases when cultures are transferred to constant temperature (**Fig 2C**). Thus, the developmental rhythms are induced by light and temperature cycles, rather than being self-sustainable.

**Figure 2.**
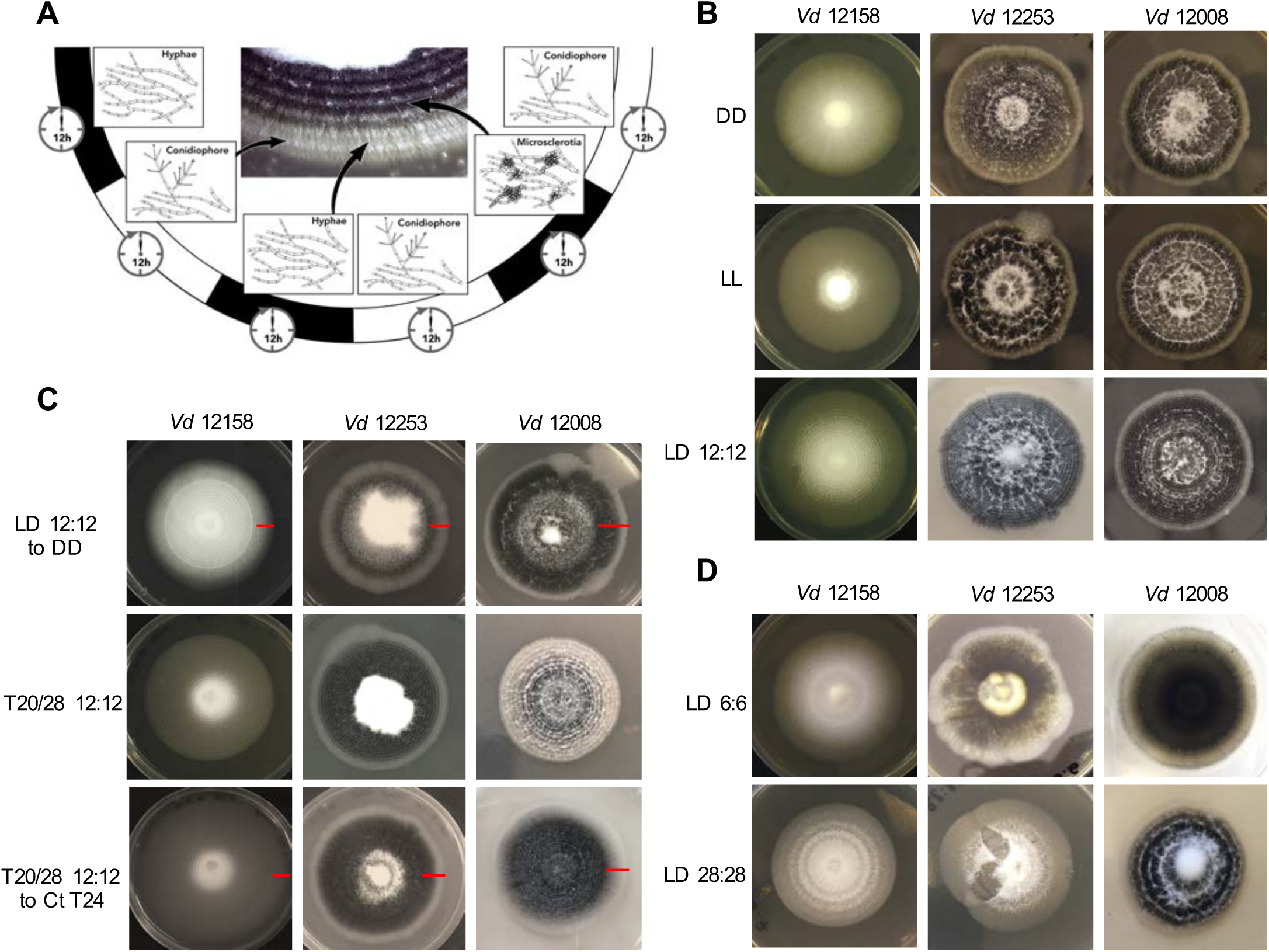
*Verticillium dahliae* morphological rhythms do not free-run. **A.** Development of *V. dahliae* in 12 h light: 12 h dark cycles (LD 12:12). After a light period, conidiophores (white bands) are developed above the hyphae, but never on the leading edge. Microsclerotia start to develop 48 h after inoculation, and are produced during the dark phase of the cycle. This leads to concentric zones of conidia and microsclerotia. **B.** *V. dahliae* morphological rhythm under different light regimes. *V. dahliae* hyaline isolate 12158 and the microsclerotia producer isolates 12253 and 12008 were point-inoculated on PLYA plates and grown in constant darkness (DD); constant white light (LL); cycles of 12 h white-light/12 h dark (LD 12:12). All plates were incubated at 24 °C. Experiments were performed five times. **C.** Strains 12158, 12253, and 12008 point-inoculated on PLYA plates and grown under light cycles of 12 h white-light:12 h dark at 24 °C or under temperature cycles of 12 h 20 °C: 12 h 28 °C (T20/28 12:12) for 14 days, were transferred to constant darkness (LD12:12-DD) or constant temperature at 24 °C (T20/28 12:12 - Ct24) for 7 days. Red horizontal lines indicate the period of growth under constant conditions. **D.** *V. dahliae* entrainment to short and long T cycles. *V. dahliae* isolates 12158, 12253 and 12008 were point inoculated onto PYLA plates and incubated for 14 days at 24 °C under T cycles of LD 6:6 and LD 28: 28.

Circadian clocks are able to entrain to cyclical cues in the environment and in all organisms studied to-date, light experienced late in the subjective day causes phase delays of clock time whilst light experienced late in the subjective night causes phase advances. This affects rhythms that are clock-controlled such that in changing day-lengths clock-controlled outputs occur at the correct time of day [3]. This ability to reset also allows circadian clocks to entrain to environmental cycles shorter and longer than 24 hours (T cycles). A phenomenon known as frequency demultiplication emerges when organisms are exposed to very short or long periods (e.g. 6:6 or 24:24 LD or temperature cycles). That is, they display a 24-hour rhythm as if the oscillation is entrained to a 12:12 cycle [3]. In contrast, if a rhythm is simply a direct response to external cues, it assumes the periodicity of the driving light or temperature cycle [3]. In *V. dahliae,* cultures grown under 6:6 LD and 6:6 temperature cycles show narrow conidial and microsclerotia bands, whereas cultures under 28:28 LD and 28:28 temperature cycles form widely spaced bands **(Fig 2D)**. Thus, the developmental rhythm shows no evidence of frequency demultiplication. Instead, conidial and microsclerotia production follows each light/dark and temperature transition.

To establish whether the lack of free-running morphological rhythms was representative of a wider set of *V. dahliae* isolates, 12 different isolates were tested under light/dark and temperature 20 °C /28 °C cycles (**S5 Fig**). Although all the isolates respond to both light and temperature cycles, in the absence of cyclic light or temperature there is no evidence of free-running rhythms. Data collected from one isolate of a species is not necessarily representative of a species [77], yet alone different species. For this reason, we characterized the photobiology and tested for the presence of circadian rhythms in three additional species in the *Verticillium* genus: *Verticillium albo-atrum*, *Verticillium nubilum* and *Verticillium tricorpus*. When cultures were treated in an alternating 12 h light/12 h dark photocycle, or 20 °C /28 °C cycles in constant darkness, concentric rings of conidia and resting structures were formed. However, as in *V. dahliae*, there is no apparent circadian rhythm of development on transfer to constant conditions in any of the Verticillium species tested in this study (**S6 Fig**).

### *V. dahliae* lacks rhythmic *Vdfrq* gene expression in LD and DD

A characteristic property of circadian rhythms is their persistence in constant conditions, and rhythmic expression of *frq* in continuous darkness has been widely used as a marker of circadian time [78]. In *V. dahliae* isolates 12253, 12008 and 12158, *Vdfrq* mRNA levels under free-running conditions are arrhythmic (*p*-value=0.94, *p*-value=1, *p*-value=0.41, respectively). In contrast, exposed to the same conditions, robust oscillation of *N. crassa frq* is seen (**Fig 3A**) and, in agreement with previous studies [79], *frq* expression peaked at 12 h and 36 h after transfer to DD (subjective morning). Another feature of circadian rhythms is that they anticipate cyclic changes in the environment. Therefore, we assessed the expression of several clock genes (*frq*, *wc-1*, *wc-2* and *vvd*) in a 12:12 LD cycle, with high time-point resolution before and after “lights on” (**Fig 3B**). *Vdfrq* transcript levels under DD seemed to anticipate the transition to light as the expression slightly increased from D10 to D11.5, albeit with less than a 1 log_2_ fold change. However, *Vdfrq* transcript levels do not exhibit significant rhythmic oscillation (*p*-value=1) and drop after the first time-point in the light. In contrast, the expression of *Vdwc-1*, *Vdwc-2* and *Vdvvd* do not anticipate the dark/light transition. There is a slight increase in the expression of *Vdwc-1* and *Vdwc-2* after 2 hours in the light, whereas *Vdvvd* is highly expressed after the dark to light transition, and drops to basal levels at a later time-points possibly due to photoadaptation [80] (**Fig 3B**). In conclusion, there is no strong evidence for anticipatory behaviour that could indicate light entrainment of a *V. dahliae* circadian clock, nor is there significant rhythmic gene expression in LD. Similarly, there is no significant difference in the expression of *Vdfrq*, *Vdwc-1*, *Vdwc-2* or *Vdccg-16* before or after the transition between a high temperature (HT) period of 28 °C or a low temperature (LT) period of 20 °C (**Fig 3C**).

**Figure 3.**
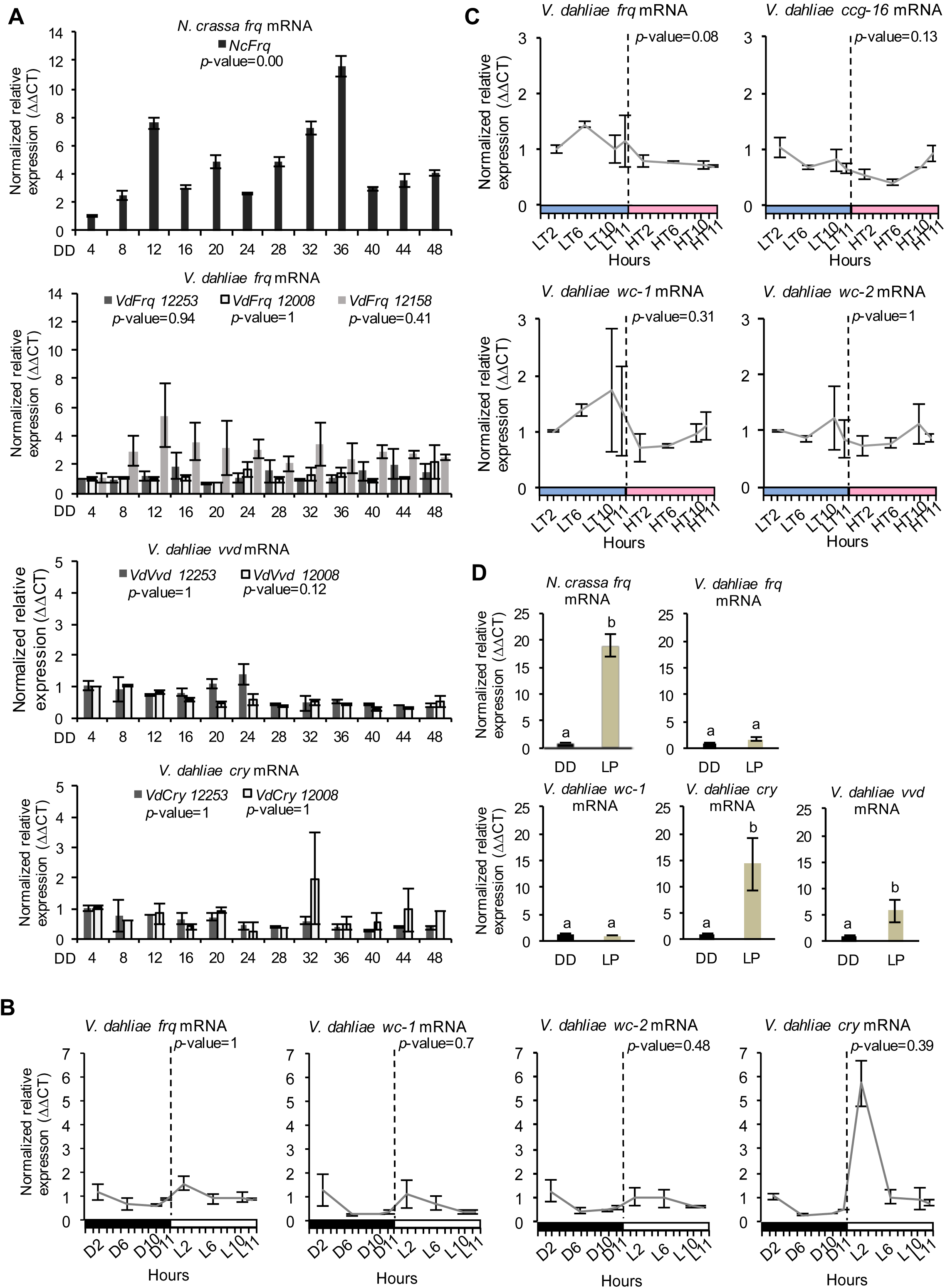
Clock-gene expression profile in *Verticillium dahliae*. **A.** Time-course expression of *N. crassa frq* and *V. dahliae frq, vvd* and *cry* under free-running conditions (DD, 24 °C) in different isolates (12253, 12008 and 12158). *VdFrq*, *Vdvvd* and *Vdcry* transcript levels were assessed by qRT-PCR every four hours over a period of 48 hours in constant darkness (DD). *N. crassa* 30-7 *bd* (first graph) was utilised as an experimental control. The β-tubulin and Elongation factor 1-⍺ genes were the *V. dahliae* housekeeping genes. The *N. crassa* housekeeping genes were the β-tubulin and TATA binding box-encoding genes. Transcript levels are normalized to ddCt of DD4 samples in each strain (control = 1). Data is presented as mean (∓SEM) from three independent biological replicates. **B.** Gene expression analysis of four *V. dahliae* putative clock genes (*Vdfrq, Vdwc-1, Vdwc-2* and *Vdvvd*). Cultures were entrained to 12 h:12 h LD cycles and *V. dahliae* 12253 tissue was harvested at different time-points in the dark (D) or light (L) over 24-hours. The dashed vertical lines represent the transition from dark to light conditions. Black-white bars indicate the dark-light conditions respectively. β-tubulin and Elongation factor 1-⍺ genes were used as housekeeping genes against which clock gene signals were normalised. Transcript levels are normalized to ddCt of D2 conditions for each gene (control = 1). Data is presented as mean (∓SEM) from three independent biological replicates. **C.** Gene expression analysis of putative clock and clock-controlled genes under (*Vdfrq, Vdwc-1, Vdwc-2* and *Vdccg-16*) cyclic temperature conditions. Cultures were entrained to 12 h/12 h 20 °C/28 °C temperature cycles, and *V. dahliae* 12253 mycelial tissues were harvested at different time-points at low (20 °C, LT) or high (28 °C, HT) temperatures over a 24-hour period. The dashed vertical lines represent the transition from L to H conditions. Blue-red bars indicate the low-high temperature conditions respectively. β-tubulin and Elongation factor 1-⍺ genes were used as housekeeping genes. Transcript levels are normalized to ddCt of L2 conditions for each gene (control = 1). Data is presented as mean (∓SEM) from three independent biological replicates. **D.** qRTPCR expression analysis of *Vdfrq*, *Vdwc-1*, *Vdcry* and *Vdvvd* in the dark and in response to a 2 minute light pulse. After 36 hours in constant dark at 25 °C, half of the cultures were given a 15 minute light pulse after which they were cultures were harvested. Statistical analysis of circadian expression was assessed by JTK-Cycle software.

### Promoter Clock box and LRE motif identification

In *N. crassa* the expression of *frq* is rapidly induced by pulses of light, mediated by the WCC, enabling light entrainment of the clock and synchronization to external conditions [5, 14]. The presence of key clock genes in Verticillium but absence of observable clock-driven developmental rhythms prompted us to look at the promoter regions of *Vdfrq* and homologues of several other genes regulated by the WCC in Neurospora. For comparison we included in our search the *frq* promoters of a subset of fungal species known to have a functional circadian clock, such as Botrytis [30] and Magnaporthe [81]. Two experimentally verified WCC binding motifs in the promoters of Neurospora clock-related genes containing two imperfect GATN repeats [7, 16], were identified in our *in silico* analysis: the proximal and distal light regulatory element (LRE) motifs containing the sequence 5’GATNC--CGATN3’, where N is the same in both repeats [45], and 5’GATCGA3’ [15].

Putative proximal or distal motifs containing the sequence 5’GATNC--CGATN3’ are found in the *frq* promoter region (2000 bp upstream the ORF) of all fungal species analysed in this study (**S7 Fig)**. However, in most species, including *V. dahlia*e, the distance between the two LRE motifs is more than 70 bp, very far from the 3 and 11 nucleotide gap found in the functionally verified proximal and distal *N. crassa frq* promoter motifs [45]. The clock-containing organism *B. cinerea* presents a putative proximal motif with a 5 nucleotide gap and *N. ditissima* shows two 4 nucleotide-gapped putative motifs within the first 400 nucleotides upstream the ORF. Interestingly, *V. tricorpus*, *V. zaregamsianum* and *V. albo-atrum* also present putative motifs with a short gap of maximum 14 nucleotides but they strongly differ in the position within the promoter. On the other hand, *Magnaporthe poae* presents a putative motif with a gap length of 55 nucleotides.

A perfect match to the proposed WCC binding motif, 5’GATCGA3’, described by Smith et al., [15] was found several times in the majority of the promoters. In *N. crassa*, the motif is found in the antisense sequence and close to the gene start site, which is also observed in *M. poae, V. nonalfalfae, V. alfalfae, V. dahliae* and *V. albo-atrum*. At the terminator level, *N. crassa* presents a WCC binding motif that induces the transcription of an antisense *frq* sequence known as *qrf* [82]. In our study, only *N. ditissima* and *V. isaaci* show a gapped motif in the antisense sequence, although they are considerably distant from the stop codon. Overall, our analysis shows that the conservation in number, position and sequence of promoter motifs does not necessarily correlate with the presence of an active circadian clock.

### Light does not activate the transcription of *Vdfrq* but regulates expression of photoreceptors *Vdcry* and *Vdvvd*

We then tested the ability of light to induce expression of *frq* in *V. dahliae*. The mycelium of *V. dahliae* 12253 strain was grown on shaking liquid media in constant darkness (DD). Then, the cultures were either kept in the dark (dark control) or transferred to white light for 15 minutes after which RNA levels were assessed by qRT-PCR (see methods). Our results reveal that *V. dahliae frq* expression is not induced by pulses of light (**Fig 3D**), whereas, as expected, *N. crassa frq* transcript increases 10-fold. Furthermore, in contrast to *N. crassa* [74], the expression of *Vdwc-1* also lacks induction by light **(Fig 3D)**. In contrast, orthologues of the light-inducible *N. crassa* genes *vvd* and *cry* are both up-regulated. *Vdcry* mRNA increases 14-fold and *Vdvvd* mRNA 5-fold in response to light **(Fig 3D)**.

### Time-course RNA-Seq reveals lack of transcriptional rhythmicity in constant conditions

Because *V. dahliae* clock gene homologues show no evidence of circadian regulation we undertook an unbiased approach to identify rhythmically expressed genes. Following light entrainment and subsequent transfer to dark, RNA-Seq analysis was carried out in three biological replicates harvested at DD6, DD12, DD18 and DD24 in wild-type (12008 and 12253) strains. The normalized data (FPKM) of each strain was analysed using JTK-Cycle [52]. The parameters were set to look for rhythms with a period of 24 hours. An important point to bear in mind is that JTK-cycle duplicates the 24-hour time series and then analyses the data for rhythmicity. Thus, as well as transcripts that reach their peak or nadir at DD12 and DD18, transcripts that decrease or increase with time after the light to dark transfer will be identified as rhythmic.

In *V. dahliae* 12253, significant oscillation in expression is observed for 568 genes (approximately 4.9 % of the genome; *p* < 0.05) although the *q* value (the minimum false discovery rate at which the test is considered significant) ranges from 0.2 to 0.8. The amplitude of the cycle is lower than 10 in 432 genes, which suggests a very weak oscillation. In *V. dahliae* 12008, 884 genes are identified with rhythmic expression, with a *p*-value < 0.05 and a *q* value ranging from 0.1 to 0.5. In this isolate *vvd, cry-dash* and putative clock-controlled genes *ccg-9* and *ccg-16* are significantly rhythmic, with amplitudes ranging from 1.4 to 21.5. However, most oscillations are weak with amplitudes lower than 10 in 83% of the rhythmically expressed genes. Of the 568 and 884 rhythmic genes identified from *V. dahliae* 12253 and 12008, only 34 genes are rhythmic in both strains **(S8 Fig)**. For those genes, the cycle amplitude is low with *q* values higher than 0.1 (**S3 Table**) and several (Chr1g02750, Chr1g18780, Chr2g02830, Chrg300650, Chr3g11370, Chr4g07950, Chr5g05660, Chr7g00930) display different expression patterns in each strain **(S8 Fig).** The expression of most genes encoding orthologues of the central clock-oscillator (*frq, wc-1, wc-2, frh, fwd-1*), clock-controlled genes (*ccg-1, ccg-6, ccg-7, ccg-8, ccg-9, ccg-14*) and photoreceptors (*phr, phy, rgs-lov, nop-1, lov-u*) are not significantly rhythmic according to the JTK Cycle analysis. Exceptions are *cry-dash* (Chr1g17210) and *vvd* (Chr3g10380) genes **(Fig 4)**. In order to validate the RNA-seq results, *V. dahliae* 12253 and 12008 samples harvested every 4 hours over a 48 hour time-course were analysed by qRT-PCR **(Fig 3A)**. In both strains, *cry-dash* and *vvd* were arrhythmic. This result highlights the possible misinterpretation of transcripts as rhythmic by JTK that are simply falling or rising after the light to dark transfer.

**Figure 4.**
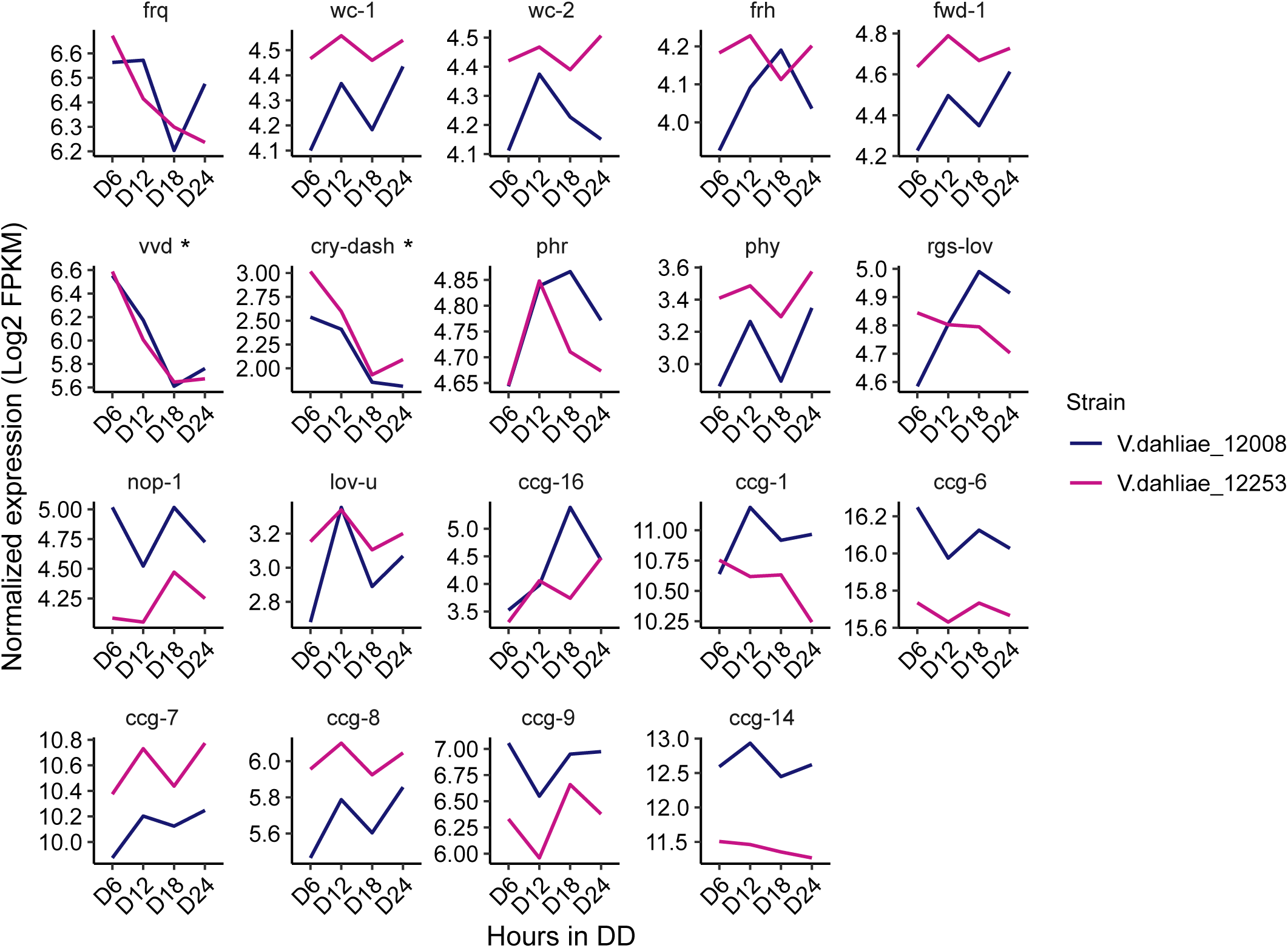
Expression pattern of *N. crassa* homologues of clock oscillator and clock-controlled genes (ccg) in both *V. dahliae* 12253 and 12008 over 24-hour in the dark. The scales are different for every gene. Statistical significant rhythmic genes assessed by JTK-Cycle are marked with an asterisk (*).

### *V. dahliae frq* affects fungal growth

Having found no strong evidence of circadian rhythmicity we wondered what function *V. dahliae* FRQ might have. To assess whether *Vdfrq* plays a role in development, we assessed whether the banding pattern of conidia and microsclerotia is generated by the *Vdfrq* knockout mutants *Δfrq_12253* and *Δfrq_12008*. The *Δfrq* mutants grown in either 12:12 h LD cycles or temperature cycles (20 °C - 28 °C) present the same characteristic banding pattern of microsclerotia and conidia as the wild type (WT) strains (**Fig 5A)**. Despite the variability among replicates we observed a reduction in total colony growth for both *Δfrq* mutants (*Δfrq_12253* and *Δfrq_12008)* with respect to the WT (12253 and 12008) strains (*p*-value < 0.01, *p*-value < 0.01, respectively). The reduction in daily growth was independent of lighting conditions (**Fig 5A**) but dependent on the nutritional composition of the culture medium (**S9 Fig**). *Δfrq_12008* showed reduced growth compared to WT_12008 on PLYA media, but not on a minimal medium (MM) (*p-*value=0.99), Czapek Dox agar (DOX) (*p*-value=1) or basal modified medium (BMM) (*p*-value=1). Colonies of *Δfrq_12253* were significantly larger than WT_12253 when incubated on BMM and MM (*p*-value < 0.01, *p*-value < 0.01, respectively). Bands of microsclerotia were observed on all media types, but conidial rings were masked by masses of mycelium when grown on DOX and BMM media (**S9 Fig**). Although a large heterogeneity exists between *Vdfrq* mutants of different *V. dahliae* isolates, the results suggest that *Vdfrq* plays a role in normal fungal growth.

**Figure 5.**
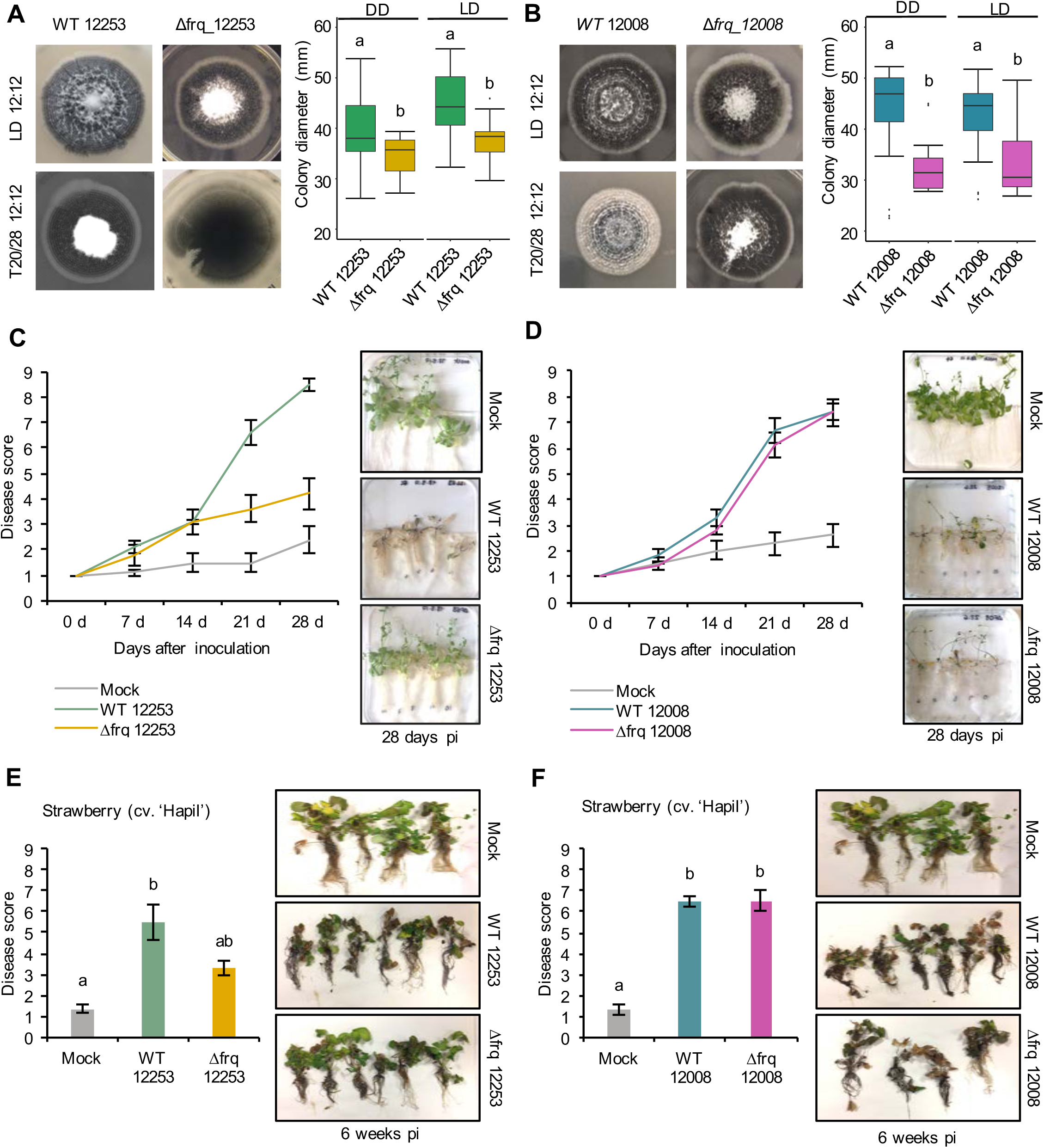
The effect of *Vdfrq* deletion mutants on fungal growth and pathogenicity. **A.** Phenotype of *V. dahliae frq* deletion in 12253 and (**B)** 12008 background. PYLA plates point inoculated with WT 12253, *Δfrq_12253,* WT 12008 and *Δfrq_12008* and incubated in LD 12:12 and temperature (20/28) 12:12 cycles for 14 days. Boxplot graphs depict WT 12253, *Δfrq_12253,* WT 12008 and *Δfrq_12008* colony diameters in DD and LD, n=6. Letters indicate statistical differences (*p*-value < 0.05), Tukey’s HSD test. **C.** Disease score of *A. thaliana in vitro* seedlings inoculated with *V. dahliae* isolates WT_12253 or *Δfrq_12253* at 0, 7, 14, 21 and 28 days post-inoculation under 12:12 LD cycle. The area under the disease progression curve (AUDPC) was calculated. Different letters indicate statistical differences (*p*-value < 0.05), Tukey’s HSD test. Disease symptoms 28 days after inoculation of *A. thaliana* seedlings with *V. dahliae* isolates and a mock (control) inoculation are shown on the right. **D.** Disease score of *A. thaliana in vitro* seedlings inoculated with *V. dahliae* isolates WT_12008 or *Δfrq_12008.* **E.** Disease score of the susceptible strawberry cultivar Hapil inoculated with WT_12253 and *Δfrq_12253,* 6 weeks after inoculation. A one-way ANOVA was performed. Data are presented as mean (∓SEM) with different letters indicating statistical differences (*p*-value < 0.05). Disease symptoms 6 weeks after inoculation of strawberry plantlets with *V. dahliae* isolates and a mock (control) inoculation are shown on the right. **F.** Disease score of the susceptible strawberry cultivar Hapil inoculated with WT 12008 and *Δfrq_12008*.

### *V. dahliae Δfrq* leads to widespread transcriptional differences

Because LREs are present in the *Vdfrq* promoter and to uncover the possible function of *Vdfrq* we compared gene expression in dark-grown WT and *Δfrq* cultures after exposure to 6 hours of white light. Of 11388 genes with nonzero total counts, 235 genes are up-regulated and 162 genes are down-regulated in the WT strain (L vs D in V.d 12253). In the *Δfrq* mutant, 65 genes are up-regulated and 30 genes are down-regulated (L vs D in *Δfrq_12253*). It is interesting that in the *Δfrq* strain, light has a reduced effect on induction/repression of gene expression suggesting that VdFRQ may indirectly affect light signaling.

The deletion of *Vdfrq* has a large effect on gene expression. Interestingly, the difference in gene expression between the *Δfrq* and the WT is greater in the light than in the dark: 278 genes are up-regulated and 195 genes are down-regulated in the *Δfrq_12253* mutant grown under dark conditions (*Δfrq* vs WT in D), whereas 435 genes are up-regulated and 233 genes are down-regulated in the *Δfrq_12253* mutants harvested after 6 h in the light (*Δfrq* vs WT in L).

Functional enrichment analysis was carried out to investigate the function of genes differentially expressed between the WT and the *Δfrq_12253* mutant. Regardless of light conditions *Δfrq_12253* up-regulated genes are involved in metabolic processes, translation, protein secretion and nucleotide metabolic processes. The down-regulated genes in *Δfrq*_12253 grown in either light or dark were involved in redox processes, heme oxidation, circadian rhythms and glutamate biosynthetic processes. Additionally, genes that are down-regulated only in the light are functionally enriched for phosphate ion transport, nitrate assimilation transport, superoxide anion generation, one-carbon metabolic processes as well as pathogenesis processes (**S4 Table**).

### *V. dahliae frq* regulates the expression of photoreceptor, TF- and SM-encoding genes

In *Neurospora*, the transcription factor complex of the clock oscillator machinery activates the transcription of many other TFs which results in a gene regulatory cascade [84]. As a result, the circadian clock modulates the expression of genes involved in many processes, especially those involved in physiology and metabolism [21]. Therefore, we hypothesized that disruption of *Vdfrq* would affect the expression of photoreceptors, transcription factor-encoding genes and genes involved in secondary metabolism.

We looked at how the absence of VdFRQ affects expression of clock, photoreceptor, transcription factor and secondary metabolism-encoding genes in *V. dahliae* under different light conditions (6 h dark and 6 h light). The expression of *fwd-1*, *wc-1* and *wc-2* are significantly down-regulated in *Δfrq_12253* in both light and dark conditions, although the LFC in expression does not reach the threshold of 1 (**S5 Table**). Interestingly, when exposed to light *vvd* is significantly up-regulated in the *Δfrq_12253* mutant, indicating that VdFRQ has a negative effect on *vvd* transcription. In the case of *cry-dash* gene, its expression is light-regulated in both the WT and *Δfrq_12253,* although the absolute expression level in the *Vdfrq* mutant is significantly lower than in the WT. The expression profile of the *rgs-lov, cry-1, phy, phr* and *nop-1* genes does not change in the absence of *Vdfrq*.

We found that the absence of *Vdfrq* also affects the expression of several TF-encoding genes. A total of 50 genes containing functional annotations associated with TF exhibited differences in expression due to the lighting conditions or strain background. Most of the differences in expression were due to the *Vdfrq* mutation. 26 TF-encoding genes were up-regulated, and 8 TF-encoding genes were down-regulated in *Δfrq_12253* regardless of the light conditions. However, there were 6 TF-encoding genes that were no longer light-induced or light-repressed in *Δfrq_12253* mutant strain (**S5 Table**).

Deletion of *Vdfrq* also affected core secondary metabolite-biosynthetic gene expression. The lovastatine nonaketide synthase encoding gene (VDAG_JR2_Chr1g23880) [84] alongside 7 other members of the PKS gene cluster (cluster 14) are strongly affected in the absence of *Vdfrq*. Interestingly, the light-induced expression of these genes was compromised in *Δfrq_12253* (**S6 Table**). Additionally, 6 genes of cluster 14 are the top most down-regulated genes in *Δfrq_12253* (LFC between −1.5 to −3.4), including the lovastatine nonaketide synthase encoding gene, TOXD protein-encoding gene and a hydrolase encoding gene. An additional cluster of PKS encoding genes, the putative aflatoxin biosynthetic cluster (cluster 17), exhibit 10 up-regulated genes in the absence of *Vdfrq*. Several members of cluster 24 (PKS), where the core biosynthetic gene encodes a fatty acid synthase, also exhibit overexpression in *Δfrq_12253*. Furthermore, 4 genes of the 5 non-ribosomal peptide synthases (NRPS, cluster 78) are highly up-regulated (LFC > 2) in the absence of *Vdfrq* (**S6 Table)**. Therefore, *Vdfrq* is crucial for the regulation of expression of secondary-metabolism-encoding genes in *V. dahliae*.

### *V. dahliae Δfrq* mutants display reduced pathogenicity in a strain-dependent manner

In *Botrytis cinerea* the circadian clock regulates virulence [85] and consistent with this finding, in a small number of fungi, circadian clock mutants show altered pathogenicity. To assess whether the loss of *Vdfrq* would influence the process of infection of *V. dahliae* we evaluated pathogenicity of two wild-type isolates (*V. dahliae* 12253 and 12008) as well as *Δfrq_12253* and *Δfrq_12008* on *A. thaliana* and *Fragaria x ananassa in vitro*-grown plants. Both wild type isolates were isolated from UK strawberries and fall within the VC group subclade II-2, 12008 being a highly virulent isolate and 12253 being a moderately virulent isolate. The infected seedlings were incubated in a 12:12 LD cycle for 28 days. Symptoms were visually rated at 0, 7, 14, 21 and 28 dpi on a scale of 1 to 9, in which 1 was equal to no symptoms and 9 equalled a dead plant. In *A. thaliana* seedlings infected with the wild-type strain WT_12253 present symptoms in up to 75% of the leaves after 21 days of inoculation (**Fig 5C**). At the same time post inoculation plants infected with *Δfrq_12253* show symptoms of wilt on 20% of leaves. The difference is more obvious at 28 dpi, when most plants infected with WT_12253 are dead whilst *Δfrq_12253* infected plants display slight chlorotic symptoms in several outer leaves. The AUDPC confirmed a significant difference in pathogenicity between *Δfrq_12253* and WT_12253 strains (*p*-value < 0.01). Contrary to this observation, WT_12008 and *Δfrq_12008* do not present differences in virulence (**Fig 5D**). Similar results are obtained from pathogenicity tests on a susceptible strawberry cultivar (Hapil). After 6 weeks of inoculation, plantlets infected with the *Δfrq_12253* strain show fewer disease symptoms than the WT_12253 strain **(Fig 5E)**, whereas the WT_12008 and *Δfrq_12008* strains do not show differences in the ability to cause disease **(Fig 5F)**. These results indicate that pathogenicity of the *Vdfrq* mutant is impaired in an isolate-dependent manner, with isolates with greater degrees of pathogenicity showing non-significant differences.

## Discussion

Evidence that the outcome of a plant-pathogen interaction can depend on the time of day at which the interaction occurs [86–88], has recently put the spotlight on the study of the circadian clock in plant pathogenic fungi. There are multiple examples of important fungal species harbouring the genetic components of the circadian clock, but the role of the clock on pathogenicity is not well understood. Optimising the processes of infection to be in synchrony with a plant’s most susceptible time could be advantageous to some fungal species for more efficient infection. Similarly, understanding the daily changes in the developmental stages of pathogenic organisms could help design more precise disease control strategies in agriculture.

In agreement with the findings of Salichos and Rokas [28], core clock orthologues were found in most of the tested Sordariomycetes species, including important plant-pathogenic fungi. Homologues of clock genes were identified in all species of the *Verticillium* genus: *V. albo-atrum, V. alfalfae, V. nonalfalfae, V. dahliae, V. longisporum* subgenome D*, V. nubilum, V. tricorpus, V. isaacii, V. klebahnii* and *V. zaregamsianum.* This result contrasts with previous analysis in which loss of the *wc-2* homologue in *V. albo-atrum* was reported [28], probably due to the poor quality at this time of the publicly available *V. albo-atrum* genome. Several species of the Dothideomycetes and Leotiomycetes, such as *Blumeria graminis, Cercospora zeae-maydis, Alternaria alternata* and *Venturia ineaqualis* lack a homologue of the blue-light receptor *vvd*, but do contain homologues of the other clock components.

The *V. dahliae* core clock homologues display strong conservation at the domain level. Remarkably, *Vd*FRQ contains all the domains identified in *N. crassa* FRQ and shares similar NLS sequences required for import of the protein to the nucleus. In addition, *Vd*FRQ exhibits conservation of phosphorylation sites, crucial for regulated activity and degradation and that determine periodicity in *N. crassa* [70]. However, small changes in domains may be crucial for function. For instance, although a coil-coil domain is present in *Vd*FRQ the probability that it can form dimers is much lower than that of the *N. crassa* FRQ coil-coil domain. Without dimerisation FRQ does not interact with the WHITE COLLAR proteins and overt rhythmicity is lost [64]. Proteins forming the WCC are also conserved in *V. dahliae*, but *Vd*WC-1 lacks the C-terminal polyQ region and has lost conservation at the N-terminal. The N-terminal of WC-1 contains important domains required for protein-protein interaction and subsequent transcriptional activation in *N. crassa* [76]. Nevertheless, zinc fingers and basic regions of both WC-1 and WC-2 required for binding DNA are present and WCC phosphosites that govern circadian repression in Neurospora [70] are conserved. Thus, *V. dahliae* contains all the components required for a TTFL but changes in some important domains may compromise their ability to generate an oscillator.

In *Neurospora* the WCCs that activate transcription in response to light and rhythmically in the dark differ in composition, in the DNA motifs they bind, and in the regions of the WCC proteins required for DNA binding. The WCC that responds to light is composed of two WC-1 proteins and one WC-2 protein. The complex binds close to the transcriptional start site of *frq* and binding requires only the zinc finger and proximal basic region of WC-2. In contrast the heterodimer of WC-1 and WC-2, responsible for transcriptional activation of *frq* in the dark, binds to the clock-box over 1 kb upstream of the transcriptional start site. Binding to the DNA requires zinc fingers and basic regions of both proteins and recruitment of chromatin modifiers SWI/SNF to initiate transcription [73]. Although several putative LREs were identified in the *Vdfrq* promoter, core sequences of the clock box that have previously been found within 1-3 bp of each other, in *V. dahliae* are separated by 80 bp. The lack of a classic clock box may underlie the lack of circadian rhythmicity of *Vdfrq* expression. However, ChIP-Seq studies have revealed diverse motifs bound by light-activated WCC [15, 83] and report that the human GATA3 protein can bind palindromic GATA sites and GATA sites located on different molecules of DNA, indicating that perhaps proximity of binding motifs is not necessarily limiting.

In order to determine whether the *V. dahliae* morphological rhythm was under the control of a circadian clock, a variety of tests were performed. Although rhythms of conidiation and microsclerotia development are observed under light-dark and temperature cycles, they do not persist in the absence of external stimuli and thus lack a key characteristic of circadian rhythms [3]. This result, repeated on different media, supports the tentative conclusion that the lack of sustainable developmental rhythms in constant conditions is unlikely to be due to media composition. Possible explanations for the absence of observable clock-controlled free-running developmental rhythms in *V. dahliae* include dampening of an existing rhythm in the absence of external signals or the lack of a functional clock.

If the former is true we theorized that the existence of a circadian clock could be revealed through analysis of development under different entrainment regimens. Frequency demultiplication effects whereby clock-controlled outputs occur once every 24 h when external periods are close to half of the endogenous period (T=12) are observed in circadian rhythms [3, 32]. Exposure to long periods (T=48) have the opposite effect, resulting in a reduction in the frequency of the output to once every 24 hours [3, 89]. *V. dahliae* grown under under 6:6 LD or 28:28 LD cycles did not exhibit frequency demultiplication and produced rings of development every 12 or 56 hours, respectively.

A defining characteristic of circadian clocks is their ability to entrain to external stimuli such as light and temperature [3]. In *N. crassa*, short pulses of light trigger a rapid induction of *frq* transcription that result in the resetting of the clock [14]. WC-1 is required for photoinduction of *frq* in response to light not only in *N. crassa* [7] but in other fungal species [29–31]. However, our results show that whereas *V. dahliae* photoreceptor-encoding genes *Vdvvd* and *Vdcry-dash* rapidly respond to light*, Vdfrq* expression is not light-induced. Furthermore, *Vdfrq, Vdwc-1, Vdwc-2, Vdvvd* and *Vdccg-16* transcript levels do not show robust anticipatory behaviour nor the significant rhythmicity in light or temperature cycles seen in other fungi with circadian clocks [29–30, 90]. Constitutive expression of *Vdfrq* under cyclic environmental conditions could be a symptom of a dysfunctional FRQ-WC clock. Alternatively, if a post-transcriptional FRQ-WC clock runs in *V. dahliae* it may represent an ancestral oscillator that in some fungi has subsequently been reinforced through additional feedback regulation acting on transcription and mRNA abundance.

RT-PCR analysis of *V. dahliae* gene expression in constant darkness after light entrainment revealed no circadian oscillation of *Vdfrq* mRNA in either of the isolates tested. However, RNA-seq gene expression studies over a 24-hour period revealed putative rhythmic expression in approximately 7% of genes, yet none of these differed more than the accepted cut-off of 1.5-fold change in expression for oscillating genes [91]. A 1.5 fold change is the threshold above which JTK_cycle reliably distinguishes between rhythmic and non-rhythmic samples. The putative rhythmic genes were also not well-correlated in the two wild-type isolates and those that were significantly rhythmic in both strains mainly showed an up or down regulation trend after the light to dark transition. It is likely that higher resolution sampling over a 48 hour time-course would greatly improve the accuracy of rhythmic transcript identification [92]. In conclusion, no strong signature of rhythmic gene expression that would indicate possible regulation of mRNA levels by a circadian clock was observed in *V. dahliae*.

In order to determine the impact of *Vdfrq* on the morphology of *V. dahliae* it was deleted in two different isolates. The absence of *Vdfrq* does not lead to the abolishment of developmental rhythms but results in reduced colony growth on most media. Moreover, pathogenicity tests reveal reduced infectivity of *Vdfrq* mutants of a weakly-pathogenic isolate but normal disease progression of a highly virulent isolate. Interestingly, this observation is repeatable across plant species (*A. thaliana* and *Fragaria x ananassa*). These data suggest that the growth penalty and/or specific changes in the expression of genes unrelated to growth in the *Vdfrq* deletion mutants influence infection and disease symptoms in a strain-dependent manner. Further study of a wide range of isolates will be needed to determine if the influence *Vdfrq* has on pathogenicity is correlated with virulence of the wildtype parent. Transcriptional profiling of a *Vdfrq* knockout mutant revealed possible roles for VdFRQ in metabolic and signaling processes and in pathogenicity. This result is in agreement with the observation that circadian control has a major impact on metabolism in *N. crassa* [21]. Interestingly, the absence of *Vdfrq* has an effect on the light response. One reason for this could be that in *ΔVdfrq* expression of *Vdwc-1* is down-regulated. This prompts speculation that a *Vdwc-1* deletion would also affect pathogenicity. To summarise, our data reveal large changes in gene expression, altered growth and pathogenicity in the *Vdfrq* deletion mutant. Whether or not these phenotypes result from *Vd*FRQ functioning outwith a circadian clock cannot at present be ascertained.

With regard to the existence of a circadian clock in *V. dahliae* our results suggest three possibilities; i) the clock is absent, ii) the clock is post-transcriptional and constitutive gene expression leads to oscillation at the protein level, iii) the clock is only active during specific developmental stages and /or specific conditions e.g. the clock is activated when a host is detected and is only functional *in planta*. As *V. dahliae* infects and moves through host tissue it is likely that an ability to anticipate time-of-day changes in host immunity would be beneficial.

At least three circumstances can be envisaged where circadian rhythmicity might be absent. The first is when an organism is always ready to respond to the rhythmic environment. It has been reported that despite the presence of homologues of most clock genes in *Picea abies* (Norway Spruce) they found no evidence of circadian gene expression in constant conditions [93]. The authors note that because gymnosperms make chlorophyll in the dark the strong adaptive pressure to anticipate dawn is lacking. Indeed, there is little evidence to support circadian gene transcription/expression in gymnosperms [94–95]. Nevertheless, night break experiments indicate that a circadian rather than an hour-glass clock is used in photoperiodism [96–98]. Secondly, in an environment where the absence of a circadian clock reduces the organism’s ability to anticipate and respond to a changing environment but this has no adverse effect. In a mutant form of einkorn wheat, rhythmicity of known clock and clock-regulated genes is lost. Counter-intuitively, rather than having a detrimental effect, in certain environments this mutant is more productive and less variable than the wild-type [99]. Importantly, the presence of an alternative circadian clock running under these conditions cannot be ruled out. Thirdly, in a predominantly arrhythmic environment, for example in underground caves and burrows where changes in temperature and humidity are minimal, or during the long winter night and perpetual daylight during mid-summer at high latitudes. Some organisms living under such conditions on initial inspection have indeed shown little or no evidence of circadian rhythmicity however, when studied in more detail these early conclusions have been overturned [100].

While definitive proof of the absence of a circadian clock is difficult to obtain, the evidence for post-transcriptional clocks and the importance of post-transcriptional modification of clock proteins is abundant and strong. A classic demonstration that post-transcriptional processes can generate a circadian clock was provided by [101] who showed that cyanobacterial clock proteins KaiA, KaiB and KiaC *in vitro* in the presence of ATP exhibit cycles of phosphorylation and dephosphorylation that have a period of approximately 24 hours, are self-sustainable and temperature compensated [101–102]. Many post-transcriptional processes act on clock gene transcripts and proteins and are key to the generation of circadian rhythmicity [103]. Indeed, it is long known that rhythms persist in enucleated *Acetabularia crenulata* [104] but there are also numerous examples of rhythmically expressed animal and plant clock genes that when constitutively expressed do not ablate rhythmicity. Rather posttranscriptional mechanisms maintain rhythmic expression and activity of the clock proteins [105–106] i.e. rhythmic transcription enhances the amplitude of rhythmic post-transcriptional processing. Indeed, even some rhythms generated post-transcriptionally are not necessarily essential parts of circadian clocks. For example, in Neurospora circadian rhythms of FRQ abundance can be decoupled from its activity [107].

The results in this study demonstrate conservation of the core clock proteins between *V. dahliae* and *N. crassa*. However, rhythmic gene expression in *V. dahliae* was not detected in either LD or free-running conditions. Thus, if a circadian clock is absent in *V. dahliae* then, at least in this fungus, the other function(s) of *Vd*FRQ must require a very similar domain structure. On the other hand, if constitutive levels of mRNA give rise to a solely protein-based circadian clock in *V. dahliae* our data also indicate that rhythmic outputs are not regulated at the level of mRNA abundance. An alternative possibility is that generation of circadian rhythmicity in *V. dahliae* is conditional on specific environmental conditions. The recent characterization of a *frq*-dependent circadian oscillator in the Leotiomycetes *Botrytis cinerea* suggests that *frq* is a component of circadian oscillators in fungal groups that evolved concurrently with *N. crassa* [29]. By extrapolation, when *frq* and other key clock genes are represented in a genome the expectation is that a FRQ-WC clock is present. This is true even when no overt rhythms in behaviour or development can be detected because clock-regulated timing of cellular biochemistry can confer a competitive advantage [108–109]. In the wild *V. dahliae* microsclerotia germinate in the presence of root exudates [36] and it is possible that this signal initiates oscillations of a circadian clock. Future studies will determine whether or not a circadian clockwork emerges *in planta* and if so what advantages this confers on the *V. dahlia* infection cycle.

## Acknowledgements

The authors acknowledge funding from BBSRC BB/R00935X/1 and BB/RR008191/1

## Author Contributions statement

EC and RH designed the experiments. EC performed the experiments. EC, SC, LJ and RH analysed the data. EC, SC, LJ and RH wrote and edited the manuscript.

## Supporting information

**S1 Fig. Generation of V. dahliae Δfrq mutants. A.** Strategy scheme of the replacement of frq gene in V. dahliae. Genomic regions utilized for homologous recombination (black fragments) are shown. Black arrows symbolise primer pairs used for the validation PCR. **B.** Gel of the validation PCRs for the correct *V. dahliae* knockout transformants. **C.** Primer pairs utilized in PCR validation.

**S2 Fig. Alignment of FRQ homologous sequences from *N. crassa* and *V. dahliae* JR2.** Protein domains characterized in *N. crassa* FRQ are annotated in red: Coiled-coil domain, FCD, FFD and PEST domains. Green bar, nuclear localization signal (NLS). Blue circle, conserved serine residue at position 513 (Ser-513). Green circles, additional phosphorylation sites previously identified in *N. crassa*.

**S3 Fig. Alignment of *N. crassa* and *V. dahliae* WC-1**. Protein domains: Dark blue poly-glutamine stretch sequences. Blue, LOV domain with black box indicating the GXNCRFLQ motif. Red, PAS domains. Yellow, predicted nuclear localization signal (NLS). Green, GATA-type zinc-finger.

**S4 Fig. Alignment of *N. crassa* and *V. dahliae* WC-2**. Protein domains: Red, PAS domain. Yellow, NLS (*V.dahliae* only). Green GATA-type zinc-finger.

**S5 Fig. The morphological rhythms of 12 *V. dahliae* isolates do not free-run.** *V. dahliae* strains isolated from multiple hosts were point-inoculated on PLYA plates and incubated for 14 days in an alternating 12 h dark/ 12 h white-light cycle (12:12 LD) (row 1) or under 12 h at 20 °C/ 12 h at 28°C (12:12 20/28) (row 3). Plates were transferred to constant darkness (12:12 LD - DD) (row 2) or constant temperature (12:12 20/28 -Ct 24) (row 4) for 7 days following the initial 14 days incubation in cyclic environments.

**S6 Fig. The morphological rhythm of *Verticillium albo-atrum* isolate 11001 and 11006, *Verticillium nubilum* isolate 15001 and *Verticillium tricorpus* isolate 20001 do not free run.** The isolates were incubated on PYLA plates for 14 days under alternating 12 h white-light/12 h dark cycles (12:12 LD) (row 1) or under 12 h at 20 °C/ 12 h at 28°C (12:12 20/28) (row 3). Plates were transferred to constant darkness (12:12 LD - DD) (row 2) or constant temperature (12:12 20/28 -Ct 24) (row 4) for 7 days following the initial 14 days incubation in cyclic environments. Red lines indicate the period of growth under constant conditions.

**S7 Fig. Schematic representation of *frq* promoter and LRE motifs in *N. crassa*, *M. poae*, *Verticillium spp.* and *B. cinerea*.** The confirmed distal LRE (dLRE) and proximal LRE (pLRE) motifs in the promoter and the qrf LRE (qLRE) motif in the terminator of *N. crassa frq* gene are marked with arrows. The putative promoter motifs containing the sequence 5’GATNC--CGATN3’ He and Liu, (2005) in the promoters (2000 bp upstream the 5’ UTR) and terminator (2000 bp downstream the 3’ UTR) are shown in blue. The 5’GATCGA3’ (Smith et al., 2010) sequences are displayed in purple

**S8 Fig. Expression profile of cycling genes detected by JTK Cycle from a 24-hour time-course RNA dataset under constant dark. A.** Venn diagram of putative rhythmic genes in *V. dahliae* 12253 and 12008 detected by JTK_Cycle. Samples were collected every 6 hours over a 24 h time-course in constant darkness at 24 °C. **B.** Expression pattern of the 34 genes identified as rhythmic by JTK-Cycle in both *V. dahliae* 12253 and 12008.

**S9 Fig. *Verticillium dahliae* growth and developmental phenotypes differ under different culture medias. A.** Growth media composition affects growth rates of WT and clock mutant strains. Quantification of colony size of the strains presented in **B**. Boxplots represent distribution of colony diameters after 14 days of incubation in 12:12 LD cycles. Two independent experiments, each containing three replicates. Analysis of variances (ANOVA) was performed. Letters indicate significant differences (*p*-value<0.05) for each media type, Tukey’s HSD test. **B.** Morphological phenotype of WT_12253, WT_12008, *Δfrq_12008, Δfrq_12253* strains were incubated on PLYA, Czapek DOX, MM and BMM plates under 12:12 LD conditions for 14 days.

**S1 Table. List of fungal strains.**

**S2 Table. List of primers.**

**S3 Table. List of rhythmically expressed genes in both *Verticillium dahliae* WT 12253, WT 12008.** Data was assessed utilising JTK-Cycle and genes with *p*-value < 0.05 are shown.

**S4 Table. List of significantly enriched GO terms related to biological processes in *Δfrq_12253* versus the WT_12253 strain in light and dark.**

**S5 Table. Expression of putative core clock genes, photoreceptor- and TF-encoding genes in WT 12253 and *Δfrq_12253* in both light and dark conditions.** Transcripts displaying a Log fold change (LFC) >1 were classified as pink, dark red if the LFC > 2, light green if the LFC < −1 and dark green if the LFC < −2. Yellow boxes indicate *p*-values < 0.05.

**S6 Table. Most differentially expressed genes in *Δfrq_12253***. Transcripts displaying a Log fold change (LFC) >1 were classified as pink, dark red if the LFC > 2, light green if the LFC < −1 and dark green if the LFC < −2. Yellow boxes indicate *p*-values < 0.05.

