## Supplementary Figures for "Homologues of key circadian clock genes present in *Verticillium dahliae* do not direct circadian programs of development or mRNA abundance"

### S1 Fig

**A**

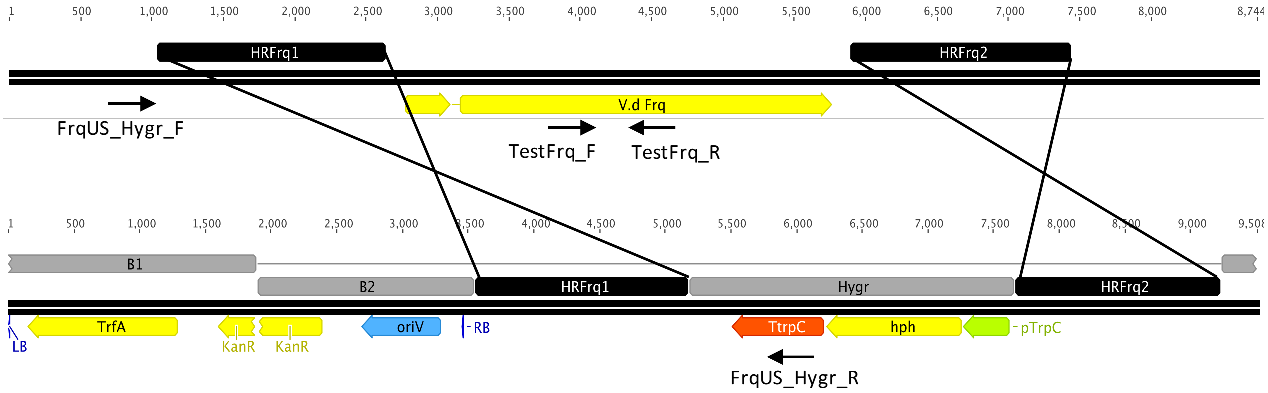


**B**

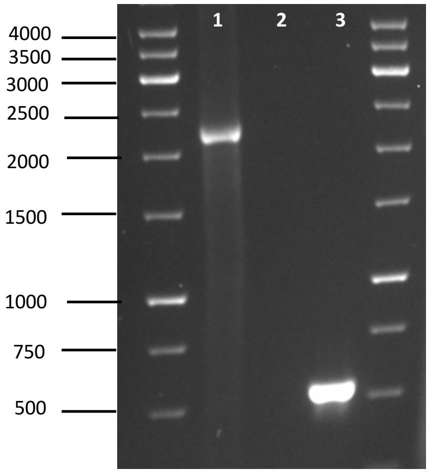


**C**

| Primer pair | Sequence 5' - 3' | Gel lane | Size (bp) |
|---|---|---|---|
| FrqUS_Hygr | AGTTCCACTCGTTCGCTCTG | 1 | 2182 |
| | CGCCTATATCGCCGACATCA | | |
| TestFrq | CCATCTTCGGCGCATTTGAG | 2, 3 | 563 |
| | ACTGTGAGGAATTGCTGCGA | | |

### S2 Fig

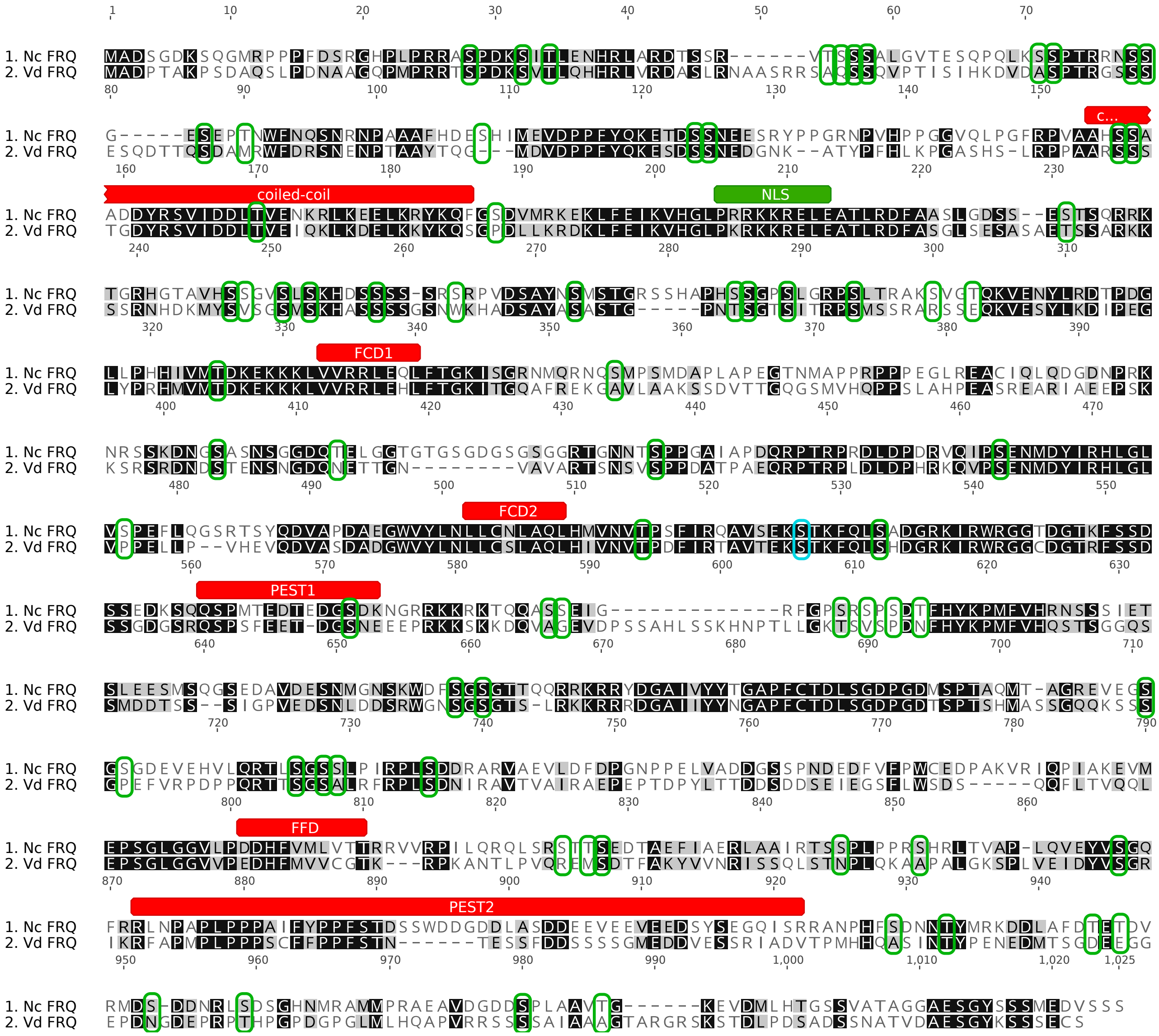

### S3 Fig

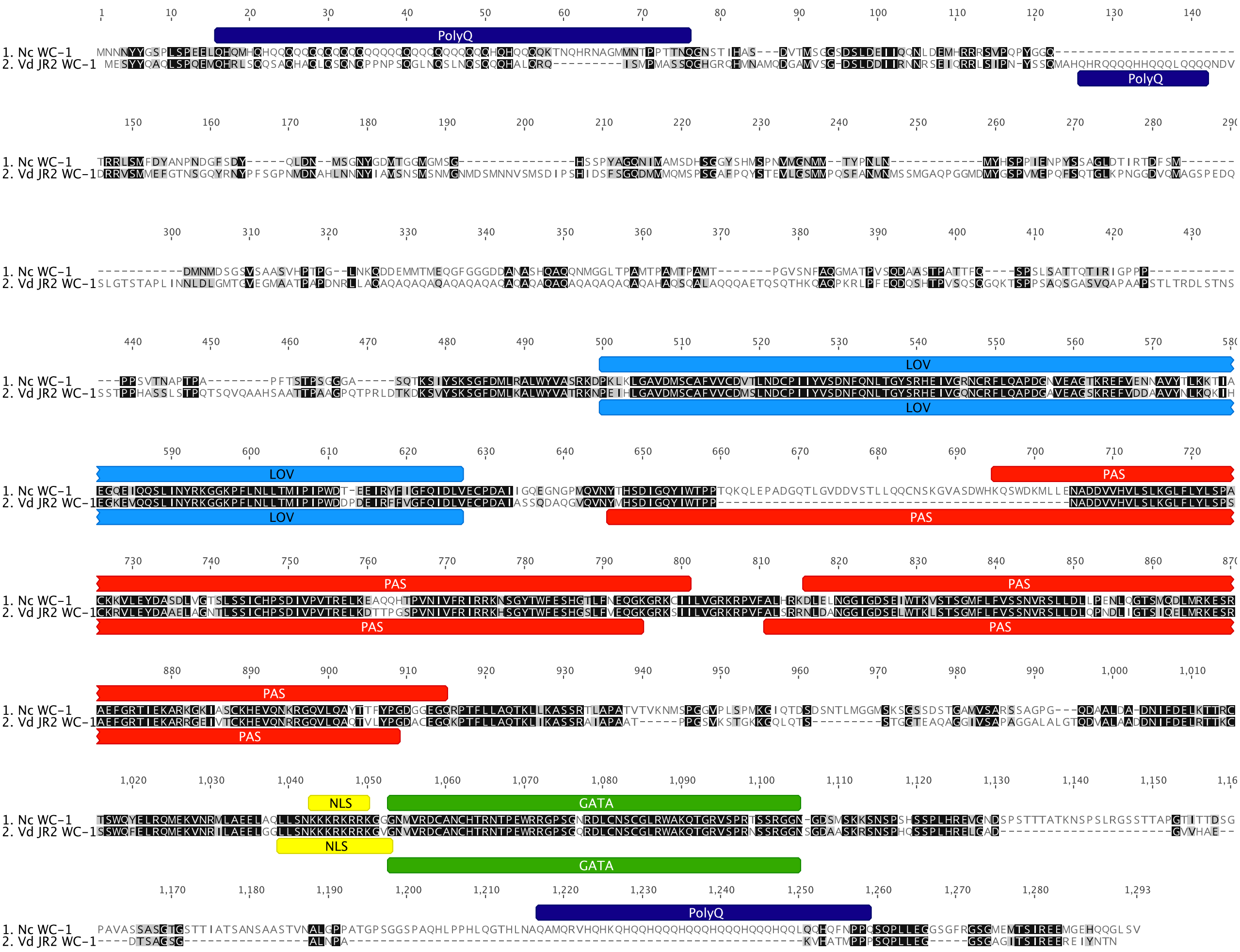

### S4 Fig

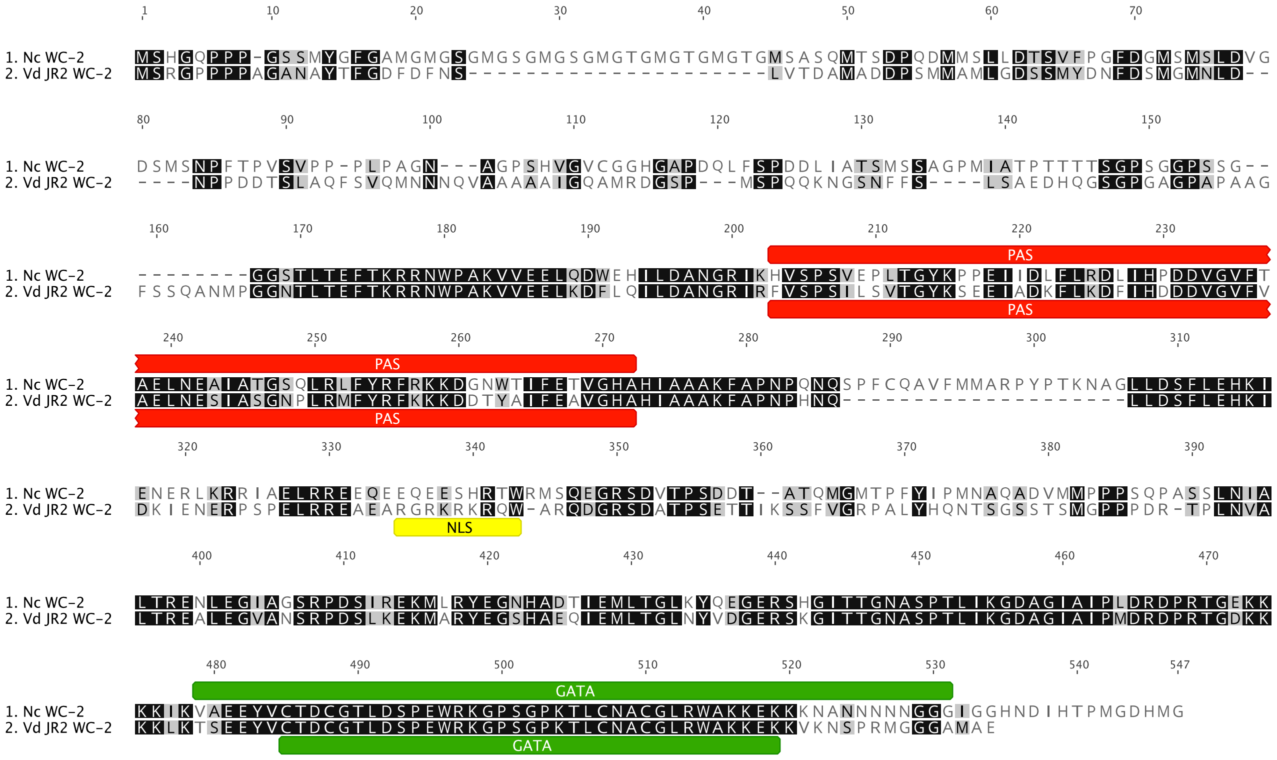

**S5 Fig**

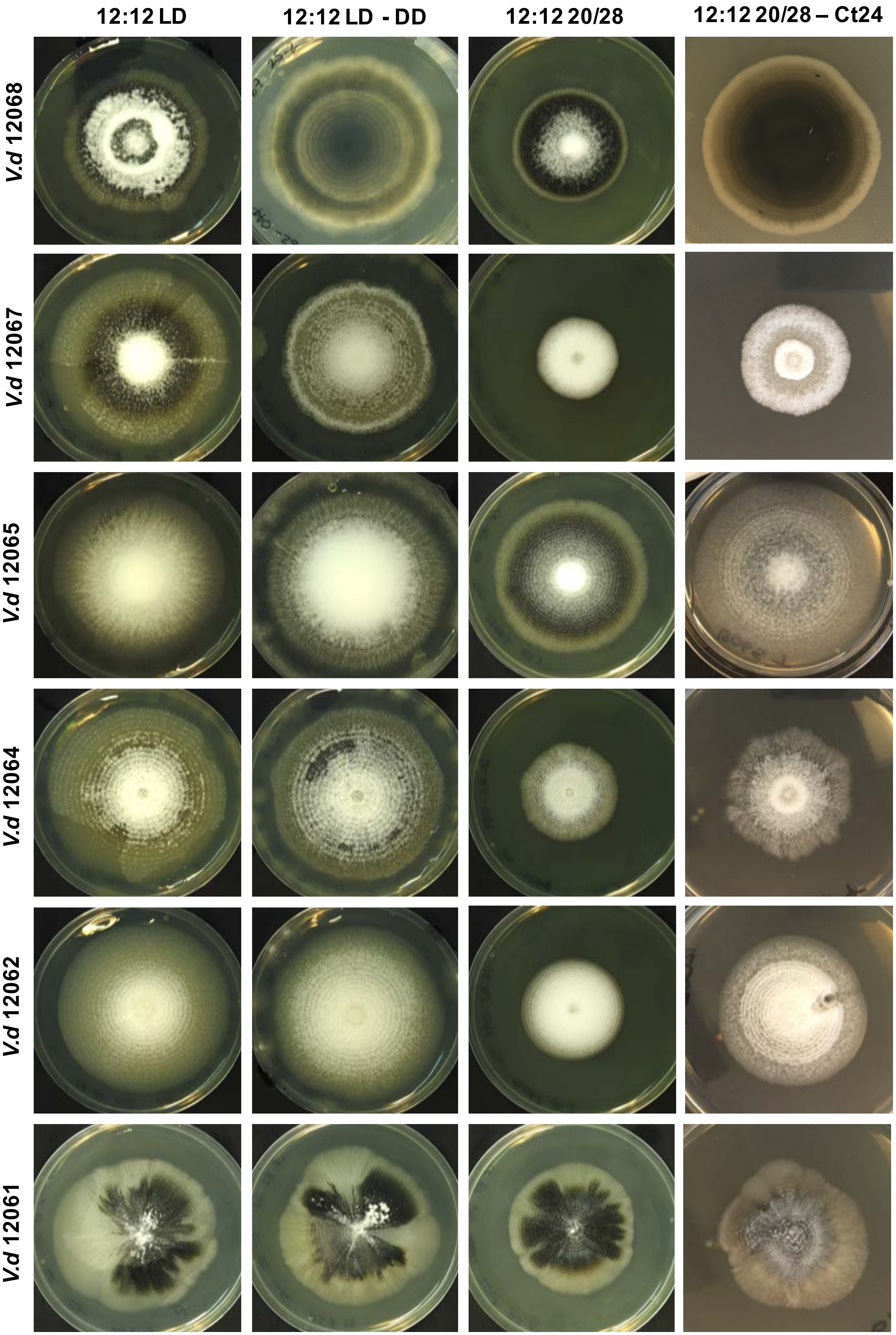
12:12 LD
12:12 LD - DD
12:12 20/28
12:12 20/28 – Ct24
*V.d* 12068
*V.d* 12067
*V.d* 12065
*V.d* 12064
*V.d* 12062
*V.d* 12061

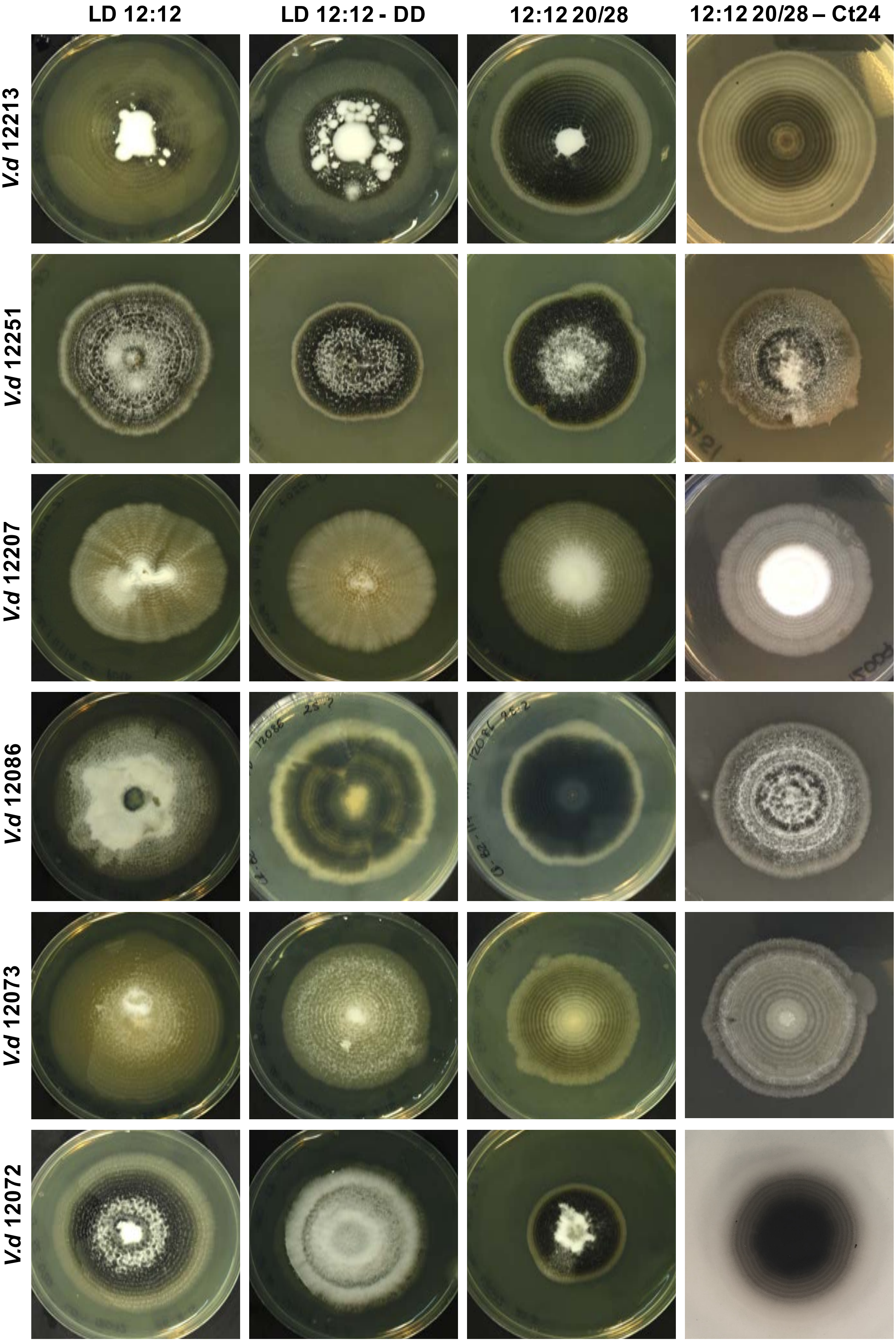

LD 12:12
LD 12:12 - DD
12:12 20/28
12:12 20/28 – Ct24
V.d 12213
V.d 12251
V.d 12207
V.d 12086
V.d 12073
V.d 12072

S6 Fig

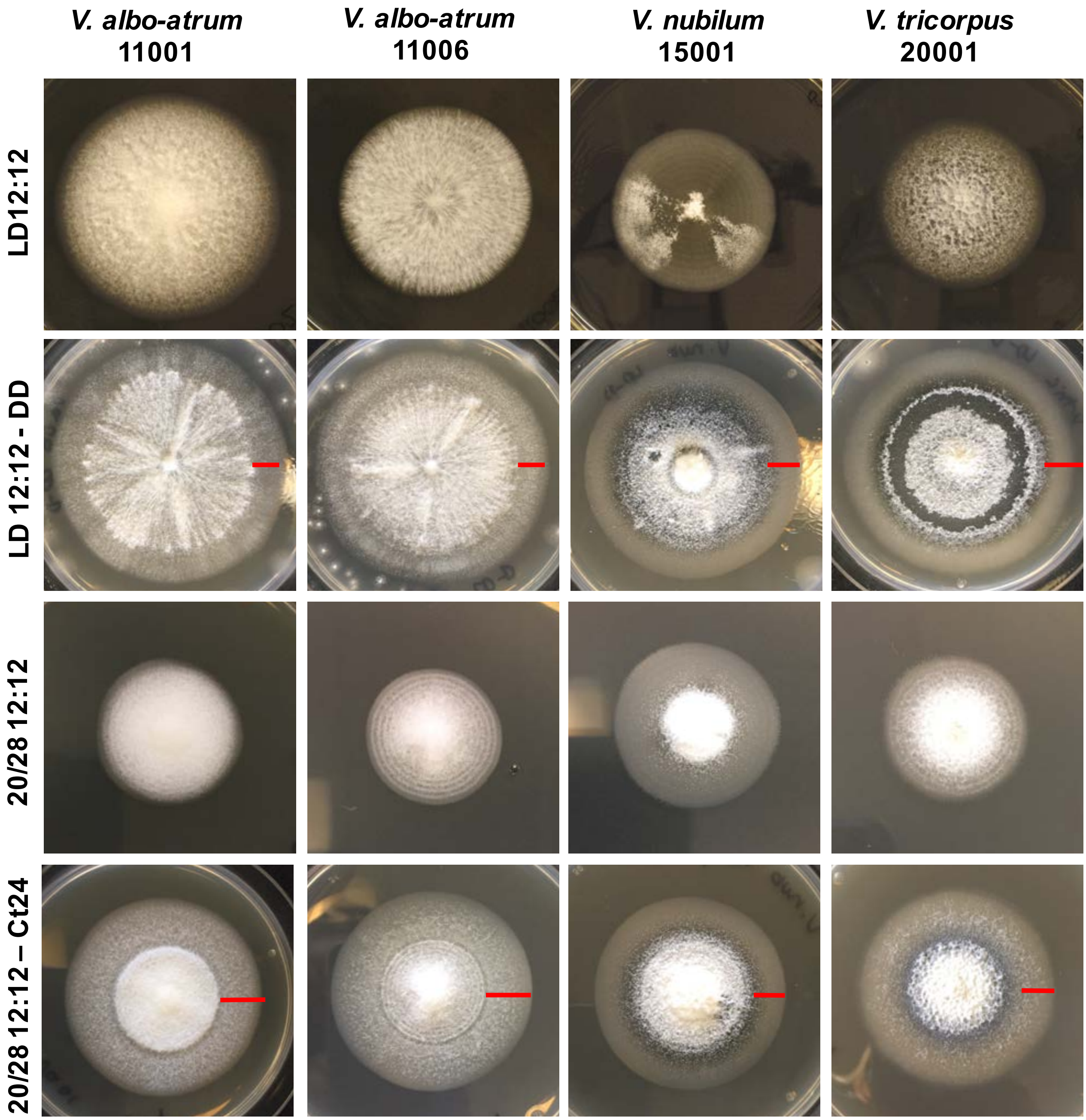
V. albo-atrum
11001
V. albo-atrum
11006
V. nubilum
15001
V. tricorpus
20001
LD12:12
LD 12:12 - DD
20/28 12:12
20/28 12:12 – Ct24

#### S7 Fig

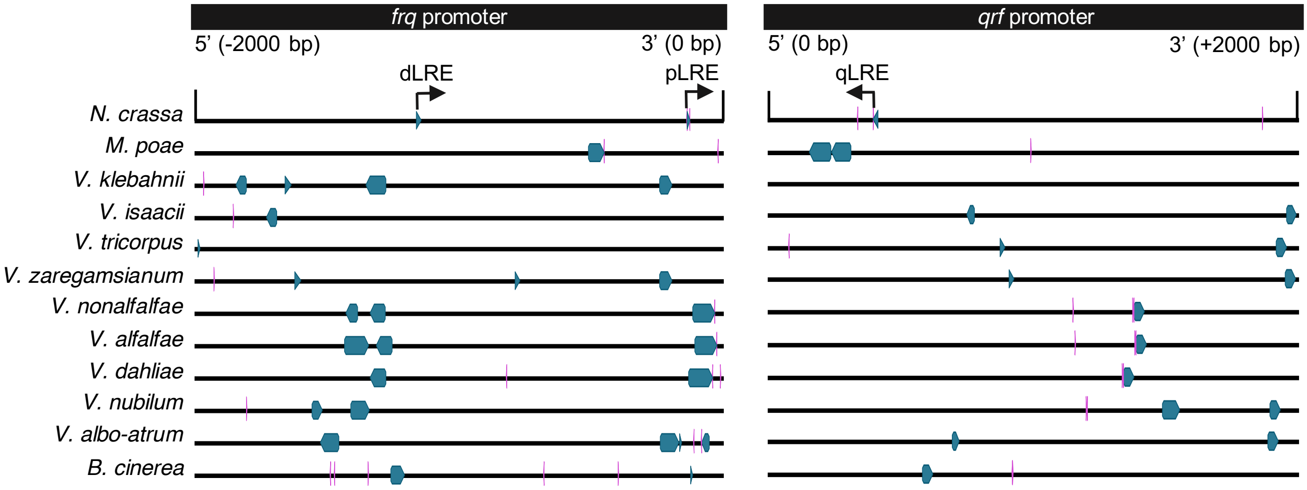
frq promoter
5' (-2000 bp)
3' (0 bp)
dLRE
pLRE
qrf promoter
5' (0 bp)
3' (+2000 bp)
qLRE
N. crassa
M. poae
V. klebahnii
V. isaacii
V. tricorpus
V. zaregamsianum
V. nonalfalfae
V. alfalfae
V. dahliae
V. nubilum
V. albo-atrum
B. cinerea

### S8 Fig

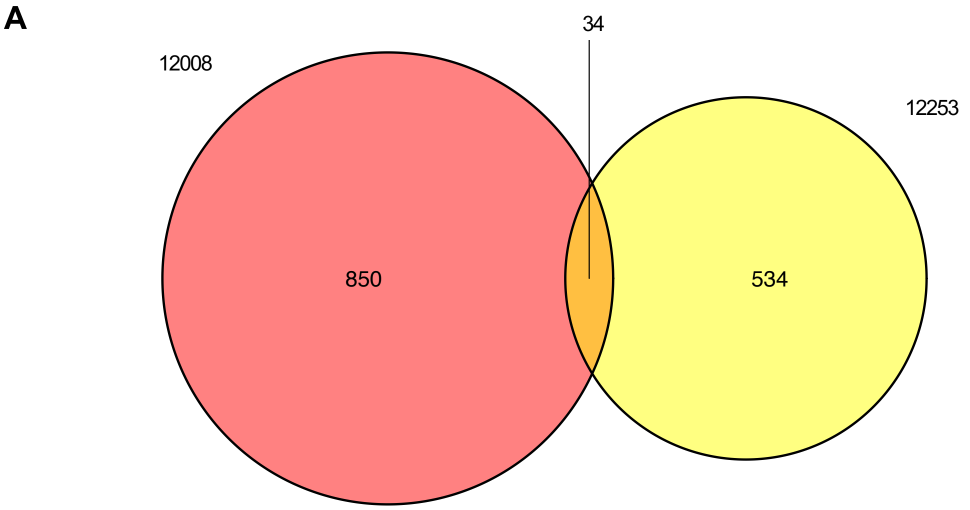
A
34
12008
12253
850
534

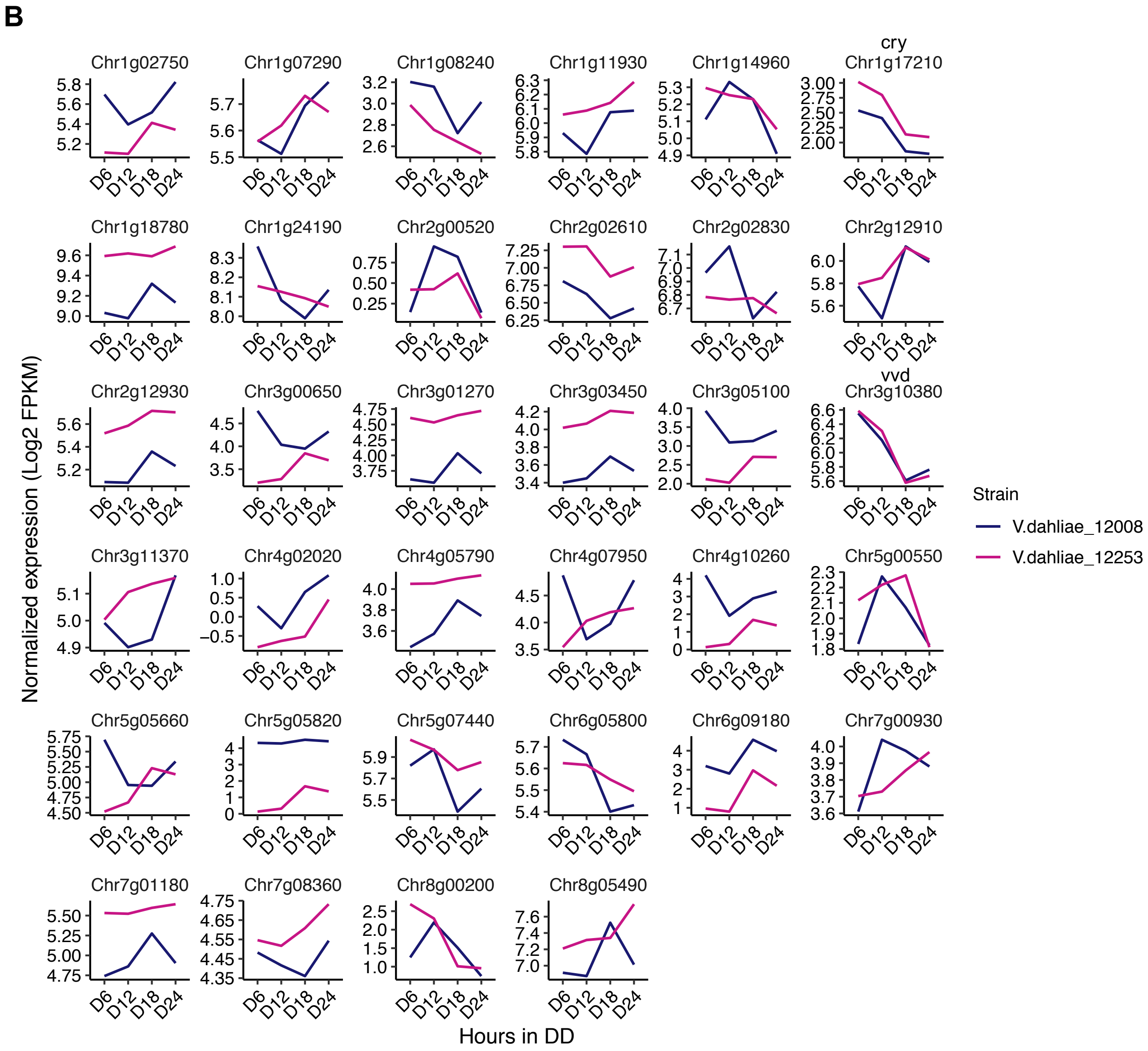
B
Chr1g02750
Chr1g07290
Chr1g08240
Chr1g11930
Chr1g14960
cry
Chr1g17210
Chr1g18780
Chr1g24190
Chr2g00520
Chr2g02610
Chr2g02830
Chr2g12910
Chr2g12930
Chr3g00650
Chr3g01270
Chr3g03450
Chr3g05100
vvd
Chr3g10380
Chr3g11370
Chr4g02020
Chr4g05790
Chr4g07950
Chr4g10260
Chr5g00550
Chr5g05660
Chr5g05820
Chr5g07440
Chr6g05800
Chr6g09180
Chr7g00930
Chr7g01180
Chr7g08360
Chr8g00200
Chr8g05490
D6
D12
D18
D24
Normalized expression (Log2 FPKM)
Hours in DD
Strain
V.dahliae_12008
V.dahliae_12253

#### S9 Fig

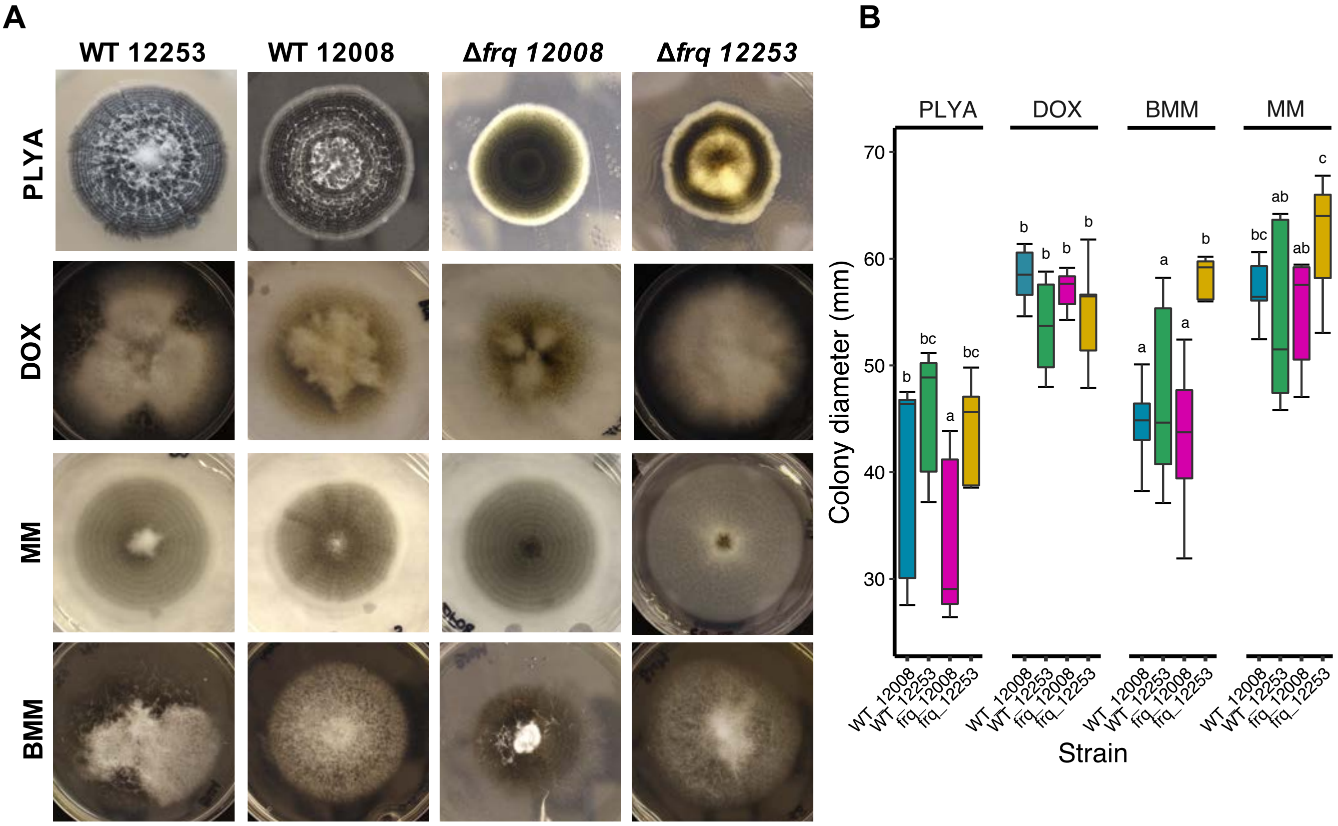

A
WT 12253
WT 12008
Δfrq 12008
Δfrq 12253
PLYA
DOX
MM
BMM
B
PLYA
DOX
BMM
MM
70
60
50
40
30
Colony diameter (mm)
b
bc
a
bc
b
b
b
b
a
b
a
a
bc
ab
ab
c
WT_12008
WT_12253
frq_12008
frq_12253
Strain
