## Supplementary Tables for "Homologues of key circadian clock genes present in *Verticillium dahliae* do not direct circadian programs of development or mRNA abundance"

**S1 Table.** List of fungal strains.

| Species | Isolate | Host | Origin | Date isolated | Source |
|---|---|---|---|---|---|
| V.dahliae | 12008 | Strawberry | Kent, UK | 1985 | Laboratory stock |
| V.dahliae | 12061 | Strawberry | Lenham, UK | 1986 | Laboratory stock |
| V.dahliae | 12062 | Strawberry | Hampshire, UK | 1965 | Laboratory stock |
| V.dahliae | 12064 | Chrysanthemum | Bristol, UK | 1968 | Laboratory stock |
| V.dahliae | 12065 | Potato | Canada, USA | 1969 | Laboratory stock |
| V.dahliae | 12067 | Tomato | Dorset, UK | 1971 | Laboratory stock |
| V.dahliae | 12068 | Hops | UK | NA | Laboratory stock |
| V.dahliae | 12072 | Gerbera | Japan | 1998 | Laboratory stock |
| V.dahliae | 12073 | Chinese Cabbage | Japan | 1998 | Laboratory stock |
| V.dahliae | 12086 | Strawberry - petiole | Kent, UK | 1998 | Laboratory stock |
| V. dahliae | 12158 | Strawberry | Kent, UK | 2000 | Laboratory stock |
| V. dahliae | 12161 | Strawberry | Lincolnshire, UK | 2000 | Laboratory stock |
| V. dahliae | 12251 | Strawberry | Kent, UK | 2012 | Laboratory stock |
| V. dahliae | 12252 | Strawberry | Kent, UK | 2012 | Laboratory stock |
| V. dahliae | 12253 | Strawberry | Kent, UK | 2012 | Laboratory stock |
| V.dahliae | 12213 | Olive | NA | 2002 | Laboratory stock |
| V. albo-atrum | 11001 | Hop | Kent, UK | 1987 | Laboratory stock |
| V. albo-atrum | 11006 | Hop | Kent, UK | 1995 | Laboratory stock |
| V. nubilum | 15001 | NA | NA | NA | Laboratory stock |
| V. tricorpus | 20001 | NA | NA | 1998 | Laboratory stock |
| V. dahliae | *Δfrq_12008* | - | - | - | This study |
| V. dahliae | *Δfrq_12253* | - | - | - | This study |
| V. dahliae | *Δwc1_12253* | - | - | - | This study |
| N. crassa | *30-7 bd a* | - | NA | NA | S. Crosthwaite |

**S2 Table.** List of primers.

| Organism | Target | Gene name | Name | Sequence (5' to 3') | Length (bp) | Sequence reference |
|---|---|---|---|---|---|---|
| V.dahliae | Housekeeping | Elongation factor alpha | EFA1-F | TACAACCCCAAGACTGTCGC | 104 | [110] |
| V.dahliae | Housekeeping | | EFA1-R | TCCTTCTCCCAGCCCTTGTA | | |
| V.dahliae | Housekeeping | Tubuline Beta Chain | VdTbc-F | TTTCCAGATCACCCACTCC | 111 | [48, 111] |
| V.dahliae | Housekeeping | | VdTbc-R | ACGACCGAGAAGGTAGCC | | |
| V.dahliae | Gene of interest | frq | VdFrq-F | GTGACGGAGAAGAGCACCAA | 92 | This study |
| V.dahliae | Gene of interest | | VdFrq-R | TCGCTACTGAATCGCGTACC | | |
| V.dahliae | Gene of interest | Wc-1 | VdWc1-F | CAACACCCTGACCGAGTTCA | 91 | This study |
| V.dahliae | Gene of interest | | VdWc1-R | ATTGGCGTCGAGAATCTGCA | | |
| V.dahliae | Gene of interest | cry | Vdcry_F | GAAGGGTGGTGAGGTGGATG | 177 | This study |
| V.dahliae | Gene of interest | | Vdcry_R | TTCACCATCTCCTCGTGCAC | | |
| V.dahliae | Gene of interest | ccg16 | Vdccg16_F | CCCAAGGCATCACAGTCGAA | 118 | This study |
| V.dahliae | Gene of interest | | Vdccg16_R | GCACCATTGTTGTTGCCGAT | | |
| V.dahliae | Gene of interest | vvd | Vdvvd_F | GATGTTCATGAACGGCGAGC | 135 | This study |
| V.dahliae | Gene of interest | | Vdvvd_R | GTCAGCCGATCTTCTCCTCG | | |
| N.crassa | Housekeeping | TATA binding box protein | NcTbp-F | GGTGCCAAGTCCGAAGATGA | 136 | [112] |
| N.crassa | Housekeeping | | NcTbp-R | TGGGGAACTTGATGTCGCAA | | |
| N.crassa | Housekeeping | Beta-tubulin | NcBtl-F | CCACTTCTTCATGGTCGGCT | 103 | [112] |
| N.crassa | Housekeeping | | NcBtl-R | CTTGGGGTCGAACATCTGCT | | |
| N.crassa | Gene of interest | Frequency | NcFrq-F | TCGACATCGCAGAGGAGAAA | 66 | [113] |
| N.crassa | Gene of interest | | NcFrq-R | CAACGAAACCCCAGACGAGT | | |

| | Primer pair | DNA Template | Overhang 5' - 3' | Sequence 5' - 3' | Length (bp) | Sequence reference |
|---|---|---|---|---|---|---|
| USER-Brick | B1-F | pRF-HU2 | AAGGTTTAAU | TCACTGGCCGTCGTTTTA | 2165 | [56] |
| | B1-R | | ATTTAAAGAU | CCGCGCGAGC | | |
| | HygR1-F | pRF-HU2 | ACGCAATACU | AGTCGGGGGATCCTCTAG | 2480 | [56] |
| | HygR1-R | | ACTAGGTCAU | GGGCCCATCGATGATCAG | | |
| | B2-F | pRF-HU2 | ATCTTTAAAU | GGAGTGTCTTCTTCCCA | 1658 | [56] |
| | B2-R | | AATACGACCU | TCGTGACTCCCTTAATTCT | | |
| | HRFrq1-F | *V. dahliae* | AGGTCGTATU | GAGACTTCAGTAAATTGTGGTTGT | 1600 | This study |
| | HRFrq1-R | | AGTATTGCGU | AGAGTCGTTGTCGTCGGG | | |
| | HRFrq2-F | *V. dahliae* | ATGACCTAGU | GATCACATTTTGCATCTTTCGGG | 1565 | This study |
| | HRFrq2-R | | ATTAAACCTU | GTCACCATGACTCGCTACAA | | |
| Plasmid validation | B1.B2-F | pEcFrq-D1 | | TGTCATACCACTTGTCCGCC | 461 | This study |
| | B1.B2-R | | | CTGCCTGTTCCAAAGGTCCT | | |
| | B1.B2.F1.H-F | pEcFrq-D1 | | ATGTTGCTGTCTCCCAGGTC | 3600 | This study |
| | B1.B2.F1.H-R | | | CCTATATCGCCGACATCACC | | |
| | H.F2.B1-F | pEcFrq-D1 | | GTATGACCGGGTCGTTCACT | 2190 | This study |
| | H.F2.B1-R | | | CTGGCTGGTGGCAGGATA | | |
| *V. dahliae* mutants validation | TestHygr_F | Δfrq mutants | | ATTTGTGTACGCCCGACAGT | 624 | This study |
| | TestHygr_R | | | AGACCTGCCTGAAACCGAAC | | |
| | TestFrq-F | Δfrq mutants | | CCATCTTCGGCGCATTTGAG | 563 | This study |
| | TestFrq-R | | | ACTGTGAGGAATTGCTGCGA | | |
| | FrqUS_Hygr-F | Δfrq mutants | | AGTTCCACTCGTTCGCTCTG | 2182 | This study |
| | FrqUS_Hygr-R | | | CGCCTATATCGCCGACATCA | | |

**S3 Table.** List of rhythmically expressed genes in both *Verticillium dahliae* WT 12253, WT 12008. Data was assessed utilising JTK-Cycle and genes with *p*-value < 0.05 are shown.

| | **V.dahliae_12253** | | | | | | | **V.dahliae_12008** | | | | | | |
|---|---|---|---|---|---|---|---|---|---|---|---|---|---|---|
| **Genes** | **q-value** | **p-value** | **Amp** | **DD6** | **DD12** | **DD18** | **DD24** | **q-Value** | **p-Value** | **Amp.** | **DD6** | **DD12** | **DD18** | **DD24** |
| VDAG_JR2_Chr3g10380 | 0.715 | 0.024 | 21.08 | 96.14 | 78.99 | 47.76 | 51.04 | 0.209 | 0.002 | 21.53 | 93.85 | 72.16 | 48.85 | 54.25 |
| VDAG_JR2_Chr4g10260 | 0.691 | 0.014 | 0.90 | 1.10 | 1.24 | 3.20 | 2.57 | 0.515 | 0.040 | 5.61 | 18.41 | 3.76 | 7.43 | 9.72 |
| VDAG_JR2_Chr5g07440 | 0.643 | 0.007 | 10.01 | 66.76 | 62.56 | 54.87 | 57.79 | 0.515 | 0.040 | 10.48 | 56.47 | 62.75 | 41.98 | 48.69 |
| VDAG_JR2_Chr1g24190 | 0.715 | 0.024 | 13.55 | 285.02 | 279.29 | 273.00 | 264.91 | 0.515 | 0.040 | 28.47 | 328.21 | 271.03 | 254.18 | 281.25 |
| VDAG_JR2_Chr1g11930 | 0.715 | 0.024 | 6.76 | 66.70 | 68.01 | 70.63 | 78.14 | 0.363 | 0.014 | 6.53 | 60.98 | 55.14 | 67.48 | 67.99 |
| VDAG_JR2_Chr3g03450 | 0.801 | 0.040 | 0.94 | 16.20 | 16.74 | 18.50 | 18.21 | 0.515 | 0.040 | 0.96 | 10.56 | 10.92 | 12.95 | 11.58 |
| VDAG_JR2_Chr4g07950 | 0.456 | 0.001 | 4.07 | 11.66 | 16.35 | 18.28 | 19.24 | 0.515 | 0.040 | 10.93 | 29.33 | 12.91 | 15.73 | 27.54 |
| VDAG_JR2_Chr6g05800 | 0.715 | 0.024 | 2.74 | 49.34 | 49.06 | 46.82 | 45.09 | 0.442 | 0.024 | 6.68 | 53.18 | 50.76 | 42.24 | 43.13 |
| VDAG_JR2_Chr5g00550 | 0.801 | 0.040 | 0.51 | 4.34 | 4.65 | 4.85 | 3.51 | 0.515 | 0.040 | 0.64 | 3.56 | 4.83 | 4.20 | 3.54 |
| VDAG_JR2_Chr2g00520 | 0.801 | 0.040 | 0.23 | 1.34 | 1.34 | 1.53 | 1.05 | 0.363 | 0.014 | 0.54 | 1.11 | 1.93 | 1.77 | 1.10 |
| VDAG_JR2_Chr4g05790 | 0.801 | 0.040 | 0.87 | 16.56 | 16.60 | 17.14 | 17.52 | 0.115 | 0.000 | 1.81 | 10.89 | 11.90 | 14.84 | 13.39 |
| VDAG_JR2_Chr5g05820 | 0.691 | 0.014 | 1.11 | 1.10 | 1.24 | 3.20 | 2.57 | 0.442 | 0.024 | 1.69 | 20.01 | 19.52 | 22.85 | 21.41 |
| VDAG_JR2_Chr8g05490 | 0.643 | 0.007 | 30.64 | 148.00 | 159.13 | 161.99 | 215.64 | 0.515 | 0.040 | 27.83 | 120.58 | 117.21 | 184.33 | 129.18 |
| VDAG_JR2_Chr7g01180 | 0.691 | 0.014 | 3.47 | 46.28 | 45.97 | 48.38 | 49.96 | 0.363 | 0.014 | 4.44 | 26.74 | 29.08 | 38.73 | 29.94 |
| VDAG_JR2_Chr1g17210 | 0.456 | 0.000 | 1.83 | 8.09 | 6.95 | 4.40 | 4.26 | 0.515 | 0.040 | 1.39 | 5.80 | 5.31 | 3.61 | 3.51 |
| VDAG_JR2_Chr1g14960 | 0.801 | 0.040 | 3.57 | 39.28 | 38.15 | 37.55 | 33.20 | 0.442 | 0.024 | 4.86 | 34.56 | 40.25 | 37.51 | 30.04 |
| VDAG_JR2_Chr7g00930 | 0.691 | 0.014 | 1.32 | 13.03 | 13.27 | 14.49 | 15.64 | 0.515 | 0.040 | 1.84 | 12.22 | 16.45 | 15.72 | 14.73 |
| VDAG_JR2_Chr8g00200 | 0.483 | 0.002 | 1.69 | 6.45 | 4.94 | 2.01 | 1.94 | 0.442 | 0.024 | 1.33 | 2.38 | 4.58 | 2.85 | 1.68 |
| VDAG_JR2_Chr2g02610 | 0.589 | 0.004 | 21.68 | 157.76 | 158.24 | 117.41 | 128.88 | 0.209 | 0.002 | 14.00 | 112.14 | 98.78 | 77.66 | 85.64 |
| VDAG_JR2_Chr4g02020 | 0.801 | 0.040 | 0.34 | 0.58 | 0.65 | 0.70 | 1.37 | 0.442 | 0.024 | 0.41 | 1.21 | 0.81 | 1.57 | 2.12 |
| VDAG_JR2_Chr3g00650 | 0.715 | 0.024 | 2.76 | 9.22 | 9.74 | 14.39 | 12.95 | 0.515 | 0.040 | 5.20 | 27.37 | 16.39 | 15.44 | 20.02 |
| VDAG_JR2_Chr6g09180 | 0.691 | 0.014 | 1.90 | 1.95 | 1.74 | 7.83 | 4.47 | 0.305 | 0.007 | 8.34 | 9.15 | 6.97 | 23.85 | 15.70 |
| VDAG_JR2_Chr2g12930 | 0.801 | 0.040 | 2.95 | 45.84 | 48.02 | 52.55 | 52.10 | 0.272 | 0.004 | 3.64 | 34.09 | 33.94 | 41.03 | 37.58 |
| VDAG_JR2_Chr1g08240 | 0.801 | 0.040 | 1.16 | 7.93 | 6.75 | 6.24 | 5.77 | 0.442 | 0.024 | 1.09 | 9.21 | 8.93 | 6.60 | 8.09 |

| | | | | | | | | | | | | | | |
|---|---|---|---|---|---|---|---|---|---|---|---|---|---|---|
| VDAG_JR2_Chr3g11370 | 0.801 | 0.040 | 1.60 | 32.08 | 34.44 | 35.17 | 35.72 | 0.305 | 0.007 | 2.49 | 31.81 | 29.89 | 30.48 | 35.95 |
| VDAG_JR2_Chr5g05660 | 0.247 | 0.000 | 8.36 | 22.93 | 25.44 | 37.54 | 34.98 | 0.442 | 0.024 | 10.65 | 51.77 | 31.00 | 30.74 | 40.45 |
| VDAG_JR2_Chr3g01270 | 0.643 | 0.007 | 2.37 | 24.36 | 23.15 | 25.08 | 26.36 | 0.515 | 0.040 | 1.84 | 12.24 | 11.78 | 16.39 | 13.09 |
| VDAG_JR2_Chr2g02830 | 0.691 | 0.014 | 7.89 | 110.27 | 108.78 | 109.65 | 101.38 | 0.442 | 0.024 | 19.46 | 124.97 | 143.06 | 98.82 | 113.23 |
| VDAG_JR2_Chr2g12910 | 0.715 | 0.024 | 8.43 | 55.45 | 57.64 | 69.57 | 64.56 | 0.305 | 0.007 | 10.10 | 54.71 | 44.87 | 70.03 | 63.57 |
| VDAG_JR2_Chr3g05100 | 0.801 | 0.040 | 2.05 | 4.35 | 4.07 | 6.55 | 6.51 | 0.166 | 0.001 | 2.64 | 15.24 | 8.53 | 8.76 | 10.58 |
| VDAG_JR2_Chr1g07290 | 0.643 | 0.007 | 2.08 | 47.12 | 49.15 | 53.12 | 50.93 | 0.115 | 0.000 | 4.78 | 47.31 | 45.64 | 51.76 | 55.08 |
| VDAG_JR2_Chr7g08360 | 0.801 | 0.040 | 1.77 | 23.37 | 22.90 | 24.39 | 26.58 | 0.442 | 0.024 | 0.91 | 22.35 | 21.35 | 20.55 | 23.32 |
| VDAG_JR2_Chr1g18780 | 0.801 | 0.040 | 32.44 | 772.28 | 786.29 | 770.98 | 824.98 | 0.515 | 0.040 | 59.47 | 523.30 | 504.24 | 639.00 | 561.44 |
| VDAG_JR2_Chr1g02750 | 0.801 | 0.040 | 3.96 | 34.64 | 34.31 | 42.58 | 40.58 | 0.363 | 0.014 | 8.04 | 51.86 | 42.11 | 45.78 | 56.57 |

**S4 Table.** List of significantly enriched GO terms related to biological processes in *Δfrq_12253* versus the WT_12253 strain in light and dark.

| | | Δfrq vs WT in D | | Δfrq vs WT in L | |
|---|---|---|---|---|---|
| **GO.ID** | **Biological process term** | **UP** | **DOWN** | **UP** | **DOWN** |
| GO:0006412 | translation | 5.40E-11 | | 2.00E-16 | |
| GO:0006414 | translational elongation | 3.40E-05 | | | |
| GO:0008152 | metabolic process | 0.022 | | 0.01 | |
| GO:0045039 | protein import into mitochondrial inner membrane | 0.024 | | | |
| GO:0009306 | protein secretion | 0.024 | | 0.032 | |
| GO:0006228 | UTP biosynthetic process | 0.024 | | 0.032 | |
| GO:0006241 | CTP biosynthetic process | 0.024 | | 0.032 | |
| GO:0046836 | glycolipid transport | 0.024 | | | |
| GO:0006183 | GTP biosynthetic process | 0.024 | | 0.032 | |
| GO:0009116 | nucleoside metabolic process | 0.042 | | | |
| GO:0033566 | gamma-tubulin complex localization | | | 0.032 | |
| GO:0055114 | oxidation-reduction process | | 2.90E-05 | | 8.20E-05 |
| GO:0007623 | circadian rhythm | | 0.016 | | 0.0221 |
| GO:0019427 | acetyl-CoA biosynthetic process from acetate | | 0.016 | | |
| GO:0006788 | heme oxidation | | 0.016 | | 0.0221 |
| GO:0009448 | gamma-aminobutyric acid metabolic processes | | 0.031 | | |
| GO:0006537 | glutamate biosynthetic process | | 0.032 | | 0.0437 |
| GO:0001682 | tRNA 5'-leader removal | | 0.032 | | |
| GO:0009450 | gamma-aminobutyric acid catabolic processes | | 0.032 | | |
| GO:0072488 | ammonium transmembrane transport | | 0.047 | | |
| GO:0006817 | phosphate ion transport | | | | 0.0046 |
| GO:0042128 | nitrate assimilation | | | | 0.0046 |
| GO:0006810 | transport | | | | 0.0129 |
| GO:0042554 | superoxide anion generation | | | | 0.0221 |
| GO:0009405 | pathogenesis | | | | 0.0221 |
| GO:0006809 | nitric oxide biosynthetic process | | | | 0.0221 |
| GO:0000160 | phosphorelay signal transduction system | | | | 0.042 |
| GO:0009395 | phospholipid catabolic process | | | | 0.0437 |
| GO:0006730 | one-carbon metabolic process | | | | 0.0437 |

**S5 Table 5.** Expression of putative core clock genes, photoreceptor- and TF-encoding genes in WT 12253 and *Δfrq_12253* in both light and dark conditions. Transcripts displaying a Log fold change (LFC) >1 were classified as pink, dark red if the LFC > 2, light green if the LFC < -1 and dark green if the LFC < -2. Yellow boxes indicate *p*- values < 0.05.

| | | L/D in V.d 12253 | | L/D in Δfrq_12253 | | Δfrq/WT in D | | Δfrq/WT in L | | | | | |
|---|---|---|---|---|---|---|---|---|---|---|---|---|---|
| **Transcript_id** | **baseMean** | **LFC** | **p-value** | **LFC** | **p-value** | **LFC** | **p-value** | **LFC** | **p-value** | **Gene** | **Antismash** | **SM** | **Interpro** |
| **Clock oscillator genes** | | | | | | | | | | | | | |
| VDAG_JR2_Chr6g03850 | 2113.89 | 0.07 | 0.51 | -0.21 | 0.10 | -0.16 | 0.08 | -0.45 | 0.00 | *fwd-1* | cluster_63 | cf_putative | F-box; WD40 repeat |
| VDAG_JR2_Chr2g01990 | 1748.38 | 0.17 | 0.23 | -0.16 | 0.47 | -0.30 | 0.02 | -0.63 | 0.00 | *wc-1* | | | Zinc finger, GATA-type |
| VDAG_JR2_Chr4g00070 | 1474.05 | -0.01 | 0.94 | -0.11 | 0.66 | 0.06 | 0.74 | -0.04 | 0.85 | *frh* | | | ATP-binding; Helicase; rRNA-processing arch, Ski2 |
| VDAG_JR2_Chr3g10380 | 1236.87 | -0.05 | 0.83 | 0.97 | 0.00 | -0.53 | 0.00 | 0.49 | 0.01 | *vvd* | | | LOV |
| VDAG_JR2_Chr7g03830 | 698.85 | -0.10 | 0.47 | -0.23 | 0.17 | -0.68 | 0.00 | -0.82 | 0.00 | *wc-2* | | | Zinc finger, GATA-type |
| **Photoreceptors genes** | | | | | | | | | | | | | |
| VDAG_JR2_Chr7g03830 | 698.85 | -0.10 | 0.47 | -0.23 | 0.17 | -0.68 | 0.00 | -0.82 | 0.00 | *wc-2* | | | Zinc finger, GATA-type |
| VDAG_JR2_Chr2g01990 | 1748.38 | 0.17 | 0.23 | -0.16 | 0.47 | -0.30 | 0.02 | -0.63 | 0.00 | *wc-1* | | | Zinc finger, GATA-type |
| VDAG_JR2_Chr3g10380 | 1236.87 | -0.05 | 0.83 | 0.97 | 0.00 | -0.53 | 0.00 | 0.49 | 0.01 | *vvd* | | | LOV |
| VDAG_JR2_Chr1g22390 | 1602.12 | 0.05 | 0.77 | -0.04 | 0.89 | 0.23 | 0.07 | 0.14 | 0.36 | *rgs-lov* | | | PAS |

| | | | | | | | | | | | | | |
|---|---|---|---|---|---|---|---|---|---|---|---|---|---|
| VDAG_JR2_Chr6g00620 | 191.19 | -0.07 | 0.82 | 0.03 | 0.95 | -0.35 | 0.09 | -0.25 | 0.29 | *cry-1* | cluster_60 | other | Photolyase/cryptochrome; FAD-binding |
| VDAG_JR2_Chr4g09150 | 1433.05 | 0.26 | 0.17 | 0.11 | 0.73 | -0.21 | 0.27 | -0.36 | 0.06 | *phy* | | | HK; PAS; Phytochrome |
| VDAG_JR2_Chr7g00170 | 1376.59 | -0.06 | 0.68 | 0.14 | 0.41 | 0.01 | 0.92 | 0.21 | 0.07 | *phr* | | | DNA-photolyase; FAD-binding |
| VDAG_JR2_Chr1g29230 | 614.73 | 0.56 | 0.18 | 0.31 | 0.63 | 0.65 | 0.09 | 0.40 | 0.36 | *nop-1* | | | G-protein glucose receptor; Rhodopsin |
| VDAG_JR2_Chr6g07410 | 386.51 | 0.56 | 0.00 | 0.19 | 0.53 | -0.40 | 0.02 | -0.77 | 0.00 | *lov-u* | | | PAS |
| VDAG_JR2_Chr1g17210 | 349.67 | 0.99 | 0.00 | 1.11 | 0.00 | -1.16 | 0.00 | -1.05 | 0.00 | *cry-dash* | cluster_8 | terpene | Photolyase/cryptochrome; FAD-binding |
| **Transcription factors** | | | | | | | | | | | | | |
| VDAG_JR2_Chr1g26850 | 91.41 | -1.03 | 0.00 | -0.19 | 0.72 | -0.93 | 0.00 | -0.09 | 0.81 | | | | Zn2 Cys6;Fungal-specific TF |
| VDAG_JR2_Chr5g05720 | 27062.18 | -1.01 | 0.00 | -0.52 | 0.07 | -0.81 | 0.00 | -0.32 | 0.19 | | | | Zinc finger, C2H2-type |
| VDAG_JR2_Chr2g04990 | 4202.92 | -1.05 | 0.00 | -0.67 | 0.06 | -0.35 | 0.20 | 0.03 | 0.93 | | | | Nucleic acid-binding proteins |
| VDAG_JR2_Chr1g24030 | 47.75 | -1.23 | 0.00 | -0.53 | 0.17 | -0.20 | 0.52 | 0.50 | 0.12 | | | | Zn2 Cys6;Fungal-specific TF |
| VDAG_JR2_Chr5g05660 | 1280.98 | 1.33 | 0.00 | 0.75 | 0.01 | -0.01 | 0.98 | -0.58 | 0.01 | | | | Zinc finger, C2H2-type |
| VDAG_JR2_Chr1g23950 | 945.69 | 1.68 | 0.00 | 0.69 | 0.24 | 0.13 | 0.81 | -0.87 | 0.05 | | cluster_14 | t1pks-nrps | Zn2 Cys6 |
| VDAG_JR2_Chr4g03300 | 951.19 | 0.51 | 0.01 | 1.09 | 0.00 | 0.35 | 0.06 | 0.93 | 0.00 | | | | Zn2 Cys6 |

| | | | | | | | | | | | | |
|---|---|---|---|---|---|---|---|---|---|---|---|---|
| VDAG_JR2_Chr6g08330 | 9249.77 | -0.76 | 0.00 | -0.45 | 0.01 | -1.27 | 0.00 | -0.95 | 0.00 | cluster_68 | cf_putative | Zinc finger, C2H2-type;Fungal TF |
| VDAG_JR2_Chr8g07070 | 412.26 | -0.40 | 0.18 | 0.18 | 0.71 | -1.13 | 0.00 | -0.55 | 0.07 | | | Zn2 Cys6 |
| VDAG_JR2_Chr3g04200 | 36396.10 | -0.91 | 0.00 | -0.38 | 0.13 | -1.11 | 0.00 | -0.58 | 0.00 | | | Zinc finger, C2H2-type |
| VDAG_JR2_Chr2g11050 | 3358.89 | 0.59 | 0.00 | 0.47 | 0.01 | 1.10 | 0.00 | 0.97 | 0.00 | | | Zn2 Cys6 |
| VDAG_JR2_Chr2g03990 | 18.20 | 0.77 | 0.11 | -0.07 | 0.94 | 1.31 | 0.00 | 0.48 | 0.32 | cluster_19 | cf_putative | Fungal-specific TF |
| VDAG_JR2_Chr8g00200 | 52.88 | -1.92 | 0.00 | 0.17 | 0.82 | -1.55 | 0.00 | 0.54 | 0.22 | | | C2H2 and C2HC zinc fingers |
| VDAG_JR2_Chr7g07100 | 23846.48 | -1.01 | 0.00 | -0.72 | 0.02 | -1.11 | 0.00 | -0.83 | 0.00 | | | HLH |
| VDAG_JR2_Chr5g05400 | 1535.28 | -0.24 | 0.12 | -0.33 | 0.11 | -0.93 | 0.00 | -1.02 | 0.00 | | | Zn2 Cys6 |
| VDAG_JR2_Chr1g03770 | 112.66 | 0.86 | 0.00 | 0.28 | 0.60 | -0.88 | 0.00 | -1.47 | 0.00 | | | Zn2/Cys6 DNA-binding domain |
| VDAG_JR2_Chr1g10470 | 1305.60 | 0.29 | 0.18 | -0.11 | 0.78 | -0.85 | 0.00 | -1.25 | 0.00 | | | Zn2 Cys6 |
| VDAG_JR2_Chr4g09890 | 2291.00 | 0.17 | 0.38 | -0.15 | 0.62 | -0.72 | 0.00 | -1.03 | 0.00 | cluster_45 | cf_putative | Zn2 Cys6 |
| VDAG_JR2_Chr3g08070 | 3513.45 | 0.75 | 0.00 | 0.29 | 0.45 | -0.67 | 0.00 | -1.13 | 0.00 | | | bZIP |
| VDAG_JR2_Chr1g27300 | 905.11 | 1.00 | 0.00 | -0.05 | 0.87 | -0.29 | 0.05 | -1.34 | 0.00 | | | Zn2 Cys6 |
| VDAG_JR2_Chr5g00990 | 503.03 | -0.58 | 0.05 | 0.24 | 0.62 | 0.34 | 0.25 | 1.16 | 0.00 | | | Zn2 Cys6;Fungal-specific TF |
| VDAG_JR2_Chr7g02150 | 631.00 | -0.30 | 0.15 | 0.37 | 0.16 | 0.43 | 0.02 | 1.10 | 0.00 | | | HLH |
| VDAG_JR2_Chr1g22480 | 307.59 | -0.20 | 0.36 | 0.30 | 0.25 | 0.56 | 0.00 | 1.06 | 0.00 | | | Zn2 Cys6 |

| | | | | | | | | | | | | |
|---|---|---|---|---|---|---|---|---|---|---|---|---|
| VDAG_JR2_Chr2g07670 | 8272.23 | -0.23 | 0.12 | 0.24 | 0.24 | 0.59 | 0.00 | 1.05 | 0.00 | | | Nucleic acid-binding proteins |
| VDAG_JR2_Chr2g10410 | 99.24 | -0.05 | 0.89 | 0.32 | 0.29 | 0.64 | 0.00 | 1.00 | 0.00 | | | Zn2 Cys6 |
| VDAG_JR2_Chr8g08220 | 276.28 | -0.15 | 0.43 | 0.23 | 0.30 | 0.65 | 0.00 | 1.03 | 0.00 | | | Nucleic acid-binding proteins |
| VDAG_JR2_Chr2g11140 | 4420.16 | -0.41 | 0.12 | 0.16 | 0.73 | 0.72 | 0.00 | 1.28 | 0.00 | | | Zinc finger, C2H2-type |
| VDAG_JR2_Chr2g03270 | 259.52 | -0.57 | 0.08 | 0.37 | 0.44 | 0.77 | 0.01 | 1.72 | 0.00 | cluster_18 | cf_putative | Zn2 Cys6;Fungal-specific TF |
| VDAG_JR2_Chr2g12610 | 11872.65 | 0.06 | 0.83 | 0.29 | 0.34 | 0.82 | 0.00 | 1.05 | 0.00 | | | Glucocorticoid receptor-like |
| VDAG_JR2_Chr5g10370 | 2513.36 | -0.79 | 0.00 | 0.11 | 0.75 | 0.83 | 0.00 | 1.73 | 0.00 | | | bZIP |
| VDAG_JR2_Chr4g07710 | 13092.22 | 0.07 | 0.78 | 0.24 | 0.42 | 0.86 | 0.00 | 1.03 | 0.00 | | | Winged helix DNA-binding domain |
| VDAG_JR2_Chr6g06230 | 2733.17 | 0.06 | 0.76 | 0.26 | 0.23 | 0.87 | 0.00 | 1.07 | 0.00 | | | Winged helix DNA-binding domain |
| VDAG_JR2_Chr4g11420 | 52.20 | -0.03 | 0.95 | 0.20 | 0.68 | 0.90 | 0.00 | 1.13 | 0.00 | cluster_48 | cf_putative | Zn2 Cys6 |
| VDAG_JR2_Chr1g03460 | 762.45 | 0.14 | 0.48 | 0.30 | 0.18 | 0.93 | 0.00 | 1.09 | 0.00 | | | Homeodomain-like |
| VDAG_JR2_Chr4g10010 | 79.48 | -0.24 | 0.51 | -0.07 | 0.90 | 0.98 | 0.00 | 1.14 | 0.00 | cluster_45 | cf_putative | MADS-box |
| VDAG_JR2_Chr3g06100 | 289.98 | -0.15 | 0.68 | 0.48 | 0.18 | 0.98 | 0.00 | 1.61 | 0.00 | | | Zn2 Cys6;Fungal-specific TF |
| VDAG_JR2_Chr8g01630 | 439.62 | -1.26 | 0.00 | 0.42 | 0.31 | -0.59 | 0.03 | 1.09 | 0.00 | | | Zinc finger, C2H2-type;Fungal TF |
| VDAG_JR2_Chr5g01890 | 247.82 | 1.56 | 0.00 | -0.16 | 0.75 | -0.49 | 0.07 | -2.22 | 0.00 | | | Zn2 Cys6;Fungal-specific TF |

| | | | | | | | | | | | | |
|---|---|---|---|---|---|---|---|---|---|---|---|---|
| VDAG_JR2_Chr5g10380 | 25.61 | -0.26 | 0.62 | 1.14 | 0.01 | 0.67 | 0.08 | 2.07 | 0.00 | | | Zn2 Cys6 |
| VDAG_JR2_Chr2g13070 | 264.38 | 0.00 | 0.99 | -0.12 | 0.77 | -1.53 | 0.00 | -1.66 | 0.00 | cluster_27 | cf_putative | Zn2/Cys6 |
| VDAG_JR2_Chr6g07650 | 2339.84 | -0.12 | 0.56 | -0.20 | 0.47 | -1.13 | 0.00 | -1.21 | 0.00 | | | C2H2 |
| VDAG_JR2_Chr3g07150 | 3982.06 | -0.15 | 0.44 | -0.20 | 0.47 | -1.02 | 0.00 | -1.06 | 0.00 | | | Homeodomain-like |
| VDAG_JR2_Chr1g05850 | 11542.15 | 0.12 | 0.61 | 0.13 | 0.71 | 1.00 | 0.00 | 1.01 | 0.00 | | | Nucleic acid-binding proteins |
| VDAG_JR2_Chr4g03340 | 253.43 | -0.02 | 0.94 | 0.18 | 0.54 | 1.08 | 0.00 | 1.29 | 0.00 | | | Zn2 Cys6 |
| VDAG_JR2_Chr2g10850 | 51.00 | 0.16 | 0.65 | 0.27 | 0.49 | 1.19 | 0.00 | 1.30 | 0.00 | | | Zn2 Cys6 |
| VDAG_JR2_Chr2g00520 | 69.36 | -0.15 | 0.77 | 0.87 | 0.08 | 1.29 | 0.00 | 2.32 | 0.00 | cluster_17 | t1pks | Winged helix DNA-binding domain |
| VDAG_JR2_Chr8g11340 | 2413.49 | 0.04 | 0.91 | -0.11 | 0.80 | 1.54 | 0.00 | 1.40 | 0.00 | | | Zinc finger, C2H2-type |
| VDAG_JR2_Chr2g00510 | 2.81 | -0.27 | 0.65 | 0.30 | NA | 1.79 | 0.00 | 2.35 | 0.00 | cluster_17 | t1pks | Homeodomain-like |
| VDAG_JR2_Chr7g10270 | 2637.45 | 0.28 | 0.30 | 0.10 | 0.83 | 2.35 | 0.00 | 2.17 | 0.00 | cluster_78 | nrps | bZIP |
| VDAG_JR2_Chr3g13030 | 42.82 | 1.35 | 0.00 | 0.67 | 0.19 | -1.10 | 0.00 | -1.78 | 0.00 | | | Zn2 Cys6 |

**S6 Table.** Most differentially expressed genes in *Δfrq_12253*. Transcripts displaying a Log fold change (LFC) >1 were classified as pink, dark red if the LFC > 2, light green if the LFC < -1 and dark green if the LFC < -2. Yellow boxes indicate *p*- values < 0.05.

| | | L/D in V.d 12253 | | L/D in Δfrq_12253 | | Δfrq/WT in D | | Δfrq/WT in L | | | | | |
|---|---|---|---|---|---|---|---|---|---|---|---|---|---|
| **Transcript_id** | **baseMean** | **LFC** | **p-value** | **LFC** | **p-value** | **LFC** | **p-value** | **LFC** | **p-value** | **Antismash** | **SM** | **TFs** | **Interpro** |
| **Top Up-regulated genes** | | | | | | | | | | | | | |
| VDAG_JR2_Chr7g10300 | 121.36 | 0.60 | 0.10 | 0.16 | 0.69 | 3.89 | 0.00 | 3.45 | 0.00 | cluster_78 | nrps | | hypothetical protein |
| VDAG_JR2_Chr4g11240 | 55.89 | 0.05 | 0.94 | 1.20 | 0.05 | 1.98 | 0.00 | 3.13 | 0.00 | cluster_47 | t1pks | | tubulin-tyrosine ligase |
| VDAG_JR2_Chr7g10290 | 151.84 | 0.59 | 0.16 | 0.21 | 0.73 | 3.48 | 0.00 | 3.09 | 0.00 | cluster_78 | nrps | | hypothetical protein |
| VDAG_JR2_Chr7g02720 | 1334.29 | -0.08 | 0.87 | 0.53 | 0.29 | 2.38 | 0.00 | 2.99 | 0.00 | | | | Integral membrane protein |
| VDAG_JR2_Chr2g00550 | 10.31 | 0.39 | 0.51 | 1.34 | 0.02 | 1.92 | 0.00 | 2.87 | 0.00 | cluster_17 | t1pks | | FAD binding domain-containing protein |
| VDAG_JR2_Chr8g11360 | 452.42 | -0.26 | 0.47 | 0.45 | 0.29 | 2.14 | 0.00 | 2.86 | 0.00 | | | | hypothetical protein |
| VDAG_JR2_Chr2g00500 | 5.95 | 0.08 | 0.92 | 0.69 | 0.31 | 2.24 | 0.00 | 2.85 | 0.00 | cluster_17 | t1pks | | hypothetical protein |
| VDAG_JR2_Chr8g11370 | 1056.59 | -0.17 | 0.65 | 0.49 | 0.25 | 2.09 | 0.00 | 2.76 | 0.00 | | | | S-adenosyl-L-methionine-dependent methyltransferase |
| VDAG_JR2_Chr2g00490 | 58.35 | 0.63 | 0.26 | 0.87 | 0.17 | 2.43 | 0.00 | 2.67 | 0.00 | cluster_17 | t1pks | | serine 3-dehydrogenase |

| | | | | | | | | | | | | | |
|---|---|---|---|---|---|---|---|---|---|---|---|---|---|
| VDAG_JR2_Chr7g10260 | 317.52 | 0.20 | 0.48 | 0.79 | 0.00 | 1.90 | 0.00 | 2.49 | 0.00 | cluster_78 | nrps | | hypothetical protein |
| VDAG_JR2_Chr7g02710 | 2265.56 | -0.80 | 0.00 | 0.12 | 0.81 | 1.49 | 0.00 | 2.42 | 0.00 | | | | Cytochrome |
| VDAG_JR2_Chr1g24880 | 24.81 | 0.20 | 0.73 | -0.41 | 0.52 | 2.99 | 0.00 | 2.38 | 0.00 | | | | cerato-ulmin hydrophobin |
| VDAG_JR2_Chr2g00510 | 2.81 | -0.27 | 0.65 | 0.30 | NA | 1.79 | 0.00 | 2.35 | 0.00 | cluster_17 | t1pks | Homeodomain-like | Myb-transcription protein |
| VDAG_JR2_Chr1g23290 | 36.88 | -0.67 | 0.08 | -0.78 | 0.06 | 2.43 | 0.00 | 2.33 | 0.00 | cluster_13 | cf_putative | | hypothetical protein |
| VDAG_JR2_Chr2g00520 | 69.36 | -0.15 | 0.77 | 0.87 | 0.08 | 1.29 | 0.00 | 2.32 | 0.00 | cluster_17 | t1pks | Winged helix | S-adenosyl-L-methionine-dependent methyltransferase |
| VDAG_JR2_Chr5g05550 | 22.44 | -0.70 | 0.06 | 0.66 | 0.10 | 0.94 | 0.00 | 2.30 | 0.00 | | | | hypothetical protein |
| VDAG_JR2_Chr8g02470 | 55.11 | 0.16 | 0.77 | 2.07 | 0.00 | 0.38 | 0.40 | 2.28 | 0.00 | | | | mannose-6-phosphate isomerase |
| VDAG_JR2_Chr2g02910 | 43.72 | 1.45 | 0.00 | 2.67 | 0.00 | 1.01 | 0.02 | 2.22 | 0.00 | | | | hypothetical protein |
| VDAG_JR2_Chr1g27430 | 31.46 | -0.73 | 0.05 | 0.38 | 0.40 | 1.11 | 0.00 | 2.22 | 0.00 | | | | hypothetical protein |
| VDAG_JR2_Chr8g11350 | 1851.53 | -0.11 | 0.80 | 0.01 | 0.99 | 2.06 | 0.00 | 2.18 | 0.00 | | | | Cytochrome |
| VDAG_JR2_Chr7g10270 | 2637.45 | 0.28 | 0.30 | 0.10 | 0.83 | 2.35 | 0.00 | 2.17 | 0.00 | cluster_78 | nrps | bZIP | bZIP transcription factor |

| | | | | | | | | | | | | | |
|---|---|---|---|---|---|---|---|---|---|---|---|---|---|
| VDAG_JR2_Chr8g11140 | 201.79 | -0.32 | 0.39 | 0.48 | 0.28 | 1.37 | 0.00 | 2.17 | 0.00 | | | | hypothetical protein |
| VDAG_JR2_Chr7g04960 | 100.68 | -0.70 | 0.11 | -0.47 | 0.44 | 1.93 | 0.00 | 2.16 | 0.00 | | | | NADP-dependent oxidoreductase domain-containing |
| VDAG_JR2_Chr2g00540 | 97.06 | -0.13 | 0.80 | 1.13 | 0.02 | 0.87 | 0.02 | 2.14 | 0.00 | cluster_17 | t1pks | | Alpha/Beta hydrolase |
| VDAG_JR2_Chr8g09650 | 122.03 | 0.79 | 0.06 | 0.61 | 0.27 | 2.26 | 0.00 | 2.09 | 0.00 | | | | Glycoside hydrolase family |
| VDAG_JR2_Chr5g10380 | 25.61 | -0.26 | 0.62 | 1.14 | 0.01 | 0.67 | 0.08 | 2.07 | 0.00 | | | Zn2 Cys6 | Zn2-Cys6 transcription factor |
| VDAG_JR2_Chr4g03070 | 89.63 | 2.26 | 0.00 | 1.51 | 0.00 | 2.70 | 0.00 | 1.95 | 0.00 | | | | Stress-induced protein, KGG |
| VDAG_JR2_Chr6g03720 | 31.42 | 1.08 | 0.02 | 0.56 | 0.37 | 2.44 | 0.00 | 1.92 | 0.00 | cluster_63 | cf_putative | | hypothetical protein |
| VDAG_JR2_Chr1g05450 | 20.37 | 0.56 | 0.32 | -0.19 | 0.81 | 2.24 | 0.00 | 1.49 | 0.00 | cluster_3 | cf_putative | | Integral membrane protein |
| VDAG_JR2_Chr6g09180 | 468.24 | 1.65 | 0.00 | 0.21 | 0.79 | 2.01 | 0.00 | 0.57 | 0.24 | cluster_70 | cf_putative | | isoamyl alcohol oxidase |
| VDAG_JR2_Chr4g10180 | 840.55 | 0.56 | 0.33 | -1.17 | 0.05 | 2.04 | 0.00 | 0.32 | 0.58 | | | | Endonuclease/Exonuclease/phosphatase |
| **Top Down-regulated genes** | | | | | | | | | | | | | |
| VDAG_JR2_Chr1g23880 | 410.36 | 4.27 | 0.00 | 0.46 | 0.53 | 0.34 | 0.53 | -3.46 | 0.00 | cluster_14 | t1pks-nrps | | lovastatin nonaketide synthase |

| | | | | | | | | | | | | |
|---|---|---|---|---|---|---|---|---|---|---|---|---|
| VDAG_JR2_Chr1g23930 | 223.91 | 3.76 | 0.00 | 0.75 | 0.25 | -0.01 | 1.00 | -3.02 | 0.00 | cluster_14 | t1pks-nrps | TOXD protein |
| VDAG_JR2_Chr1g23890 | 459.89 | 3.58 | 0.00 | 0.60 | 0.36 | -0.01 | 0.99 | -2.99 | 0.00 | cluster_14 | t1pks-nrps | hypothetical protein |
| VDAG_JR2_Chr8g09780 | 57.25 | 1.70 | 0.00 | -0.20 | 0.81 | -1.07 | 0.01 | -2.97 | 0.00 | | | hypothetical protein |
| VDAG_JR2_Chr1g23910 | 116.12 | 4.13 | 0.00 | 1.16 | 0.05 | 0.04 | 0.95 | -2.93 | 0.00 | cluster_14 | t1pks-nrps | hypothetical protein |
| VDAG_JR2_Chr1g23920 | 114.15 | 3.40 | 0.00 | 0.47 | 0.50 | 0.08 | 0.91 | -2.86 | 0.00 | cluster_14 | t1pks-nrps | Alpha/Beta hydrolase |
| VDAG_JR2_Chr1g23940 | 625.82 | 3.36 | 0.00 | 0.69 | 0.30 | 0.10 | 0.88 | -2.58 | 0.00 | cluster_14 | t1pks-nrps | hypothetical protein |
| VDAG_JR2_Chr2g03470 | 5765.15 | 0.98 | 0.01 | -0.19 | 0.78 | -1.37 | 0.00 | -2.54 | 0.00 | | | cytochrome P450 monooxygenase |
| VDAG_JR2_Chr8g09260 | 36.24 | 1.08 | 0.02 | 0.29 | 0.71 | -1.66 | 0.00 | -2.44 | 0.00 | | | hypothetical protein |
| VDAG_JR2_Chr6g03670 | 1016.68 | -1.99 | 0.00 | -0.15 | 0.82 | -4.27 | 0.00 | -2.44 | 0.00 | cluster_63 | cf_putative | hypothetical protein |
| VDAG_JR2_Chr1g22700 | 4387.46 | 0.46 | 0.07 | -0.34 | 0.37 | -1.58 | 0.00 | -2.38 | 0.00 | | | hypothetical protein |

| | | | | | | | | | | | | | |
|---|---|---|---|---|---|---|---|---|---|---|---|---|---|
| VDAG_JR2_Chr4g04340 | 2563.03 | -1.19 | 0.00 | 0.37 | 0.55 | -3.87 | 0.00 | -2.32 | 0.00 | | | | hypothetical protein |
| VDAG_JR2_Chr6g04130 | 796.94 | 0.58 | 0.11 | 0.09 | 0.91 | -1.79 | 0.00 | -2.28 | 0.00 | | | | hypothetical protein |
| VDAG_JR2_Chr4g11840 | 41.01 | 1.10 | 0.00 | -0.01 | 0.99 | -1.15 | 0.00 | -2.27 | 0.00 | | | | hypothetical protein |
| VDAG_JR2_Chr5g01890 | 247.82 | 1.56 | 0.00 | -0.16 | 0.75 | -0.49 | 0.07 | -2.22 | 0.00 | | | Zn2/Cys6-Fungal TF | Zn2-Cys6 transcription factor |
| VDAG_JR2_Chr6g08940 | 153.16 | 1.22 | 0.01 | 0.42 | 0.56 | -1.38 | 0.00 | -2.19 | 0.00 | cluster_69 | cf_putative | | cytochrome P450 |
| VDAG_JR2_Chr8g06660 | 13626.04 | 0.00 | 0.99 | -0.68 | 0.06 | -1.45 | 0.00 | -2.14 | 0.00 | | | | Ammonium/urea transporter |
| VDAG_JR2_Chr8g09770 | 50.00 | 1.05 | 0.01 | -0.36 | 0.58 | -0.72 | 0.07 | -2.13 | 0.00 | | | | hypothetical protein |
| VDAG_JR2_Chr1g22660 | 1015.42 | 0.87 | 0.00 | 0.06 | 0.92 | -1.23 | 0.00 | -2.04 | 0.00 | | | | UDP-glucose 4-epimerase |
| VDAG_JR2_Chr1g01960 | 4976.56 | -0.01 | 0.98 | 0.77 | 0.00 | -2.82 | 0.00 | -2.04 | 0.00 | | | | Frequency clock protein |
| VDAG_JR2_Chr1g09750 | 2398.82 | 0.84 | 0.00 | -0.04 | 0.94 | -1.16 | 0.00 | -2.03 | 0.00 | | | | cholesterol oxidase |

| | | | | | | | | | | |
|---|---|---|---|---|---|---|---|---|---|---|
| VDAG_JR2_Chr6g06730 | 959.89 | 0.33 | 0.59 | -1.68 | 0.00 | 0.01 | 0.99 | -2.01 | 0.00 | Protein of unknown function |
| VDAG_JR2_Chr3g06750 | 104.25 | -1.28 | 0.01 | 0.30 | 0.71 | -2.77 | 0.00 | -1.20 | 0.01 | hypothetical protein |
| VDAG_JR2_Chr5g10060 | 42.49 | -0.46 | 0.39 | 0.17 | 0.84 | -2.30 | 0.00 | -1.67 | 0.00 | hypothetical protein |
| VDAG_JR2_Chr1g09640 | 1869.91 | -0.26 | 0.53 | 0.54 | 0.28 | -2.02 | 0.00 | -1.22 | 0.00 | Integral membrane protein |
